# Structural studies of KCTD1 and its disease-causing mutant P20S provide insights into the protein function and misfunction

**DOI:** 10.1101/2024.06.14.599007

**Authors:** Nicole Balasco, Alessia Ruggiero, Giovanni Smaldone, Giovanni Pecoraro, Luigi Coppola, Luciano Pirone, Emilia M. Pedone, Luciana Esposito, Rita Berisio, Luigi Vitagliano

## Abstract

Members of the KCTD protein family play key roles in fundamental physio-pathological processes. A plethora of literature studies have demonstrated their involvement in cancer, neurodevelopmental disorders, and genetic diseases. Despite two decades of intense investigations, the definition of structure-(mis)function relationships for these proteins is still rather limited. Here, we derived atomic-level structural data on KCTD1, by determining the crystal structure of its P20S mutant, which causes the scalp-ear-nipple syndrome, and performing molecular dynamics simulations. In addition to the expected folded domains (BTB and CTD) the crystal structure unravels that also the N-terminal region that precedes the BTB domain (preBTB) adopts a folded polyproline II (PPII) state. The global structure of the KCTD1 pentamer is characterized by an intricate architecture in which the different subunits mutually exchange domains to generate a closed domain swapping motif. In this framework, the BTB domain of each chain makes peculiar contact with the preBTB and the CTD regions of an adjacent chain. Indeed, the BTB-preBTB interaction is made of a PPII-PPII recognition motif whereas the BTB-CTD contacts are mediated by an unusual (+/-) helix discontinuous association. The inspection of the protein structure, along with the data that emerged from the MD data, provides a clear explanation of the pathogenicity of the SENS mutation P20S and unravels the role of the BTB-preBTB interaction in the insurgence of the disease. Finally, the presence of potassium bound to the central cavity of the CTD pentameric assembly provides insights into the role of the protein in metal homeostasis.

## 1. Introduction

KCTD (K-potassium channel tetramerization domain-containing proteins) constitutes a class of emerging proteins that are involved in key physiopathological processes. The common element of this protein family is the presence, at the N-terminal region of all members, of a BTB domain (1) that presents a significant similarity with the tetramerization domain of the Kv4.2 potassium channels (2). However, despite this similarity, there is no evidence that KCTD proteins, with the possible exception of KCNRG (2), may be involved in the potassium transport. An increasing number of independent studies carried out in the last decade have demonstrated that dysregulations of these proteins have been detected in the etiology of many diversified pathological states, including cancer (3), neurodevelopmental/neuropsychiatric (4), and genetic diseases (5–11). For many years, a full understanding of the functionality of these proteins has been hampered by the paucity of related atomic-level structural data (12–14). This scenario has rapidly changed in the last few years through the combination of experimental structural studies (15–20) and approaches based on machine-learning techniques (21–23). These studies have provided a plausible global view of the structural features of these proteins and their interaction with some important biological partners. Based on these predictive studies, it has been shown that even the C-terminal domains (CTD) of these proteins, which were believed to be independent and evolutionary distinct modules, share some significant structural similarities throughout the family (21). Structure-based clustering based on the similarity of the CTD domains has highlighted the segregation of two groups of the family (cluster 1A - KCTD8/KCTD12/KCTD16 and cluster 1B - KCTD1/KCTD15) that are not involved in protein ubiquitination/degradation. Indeed, members of these two clusters do not bind Cullin 3 (CUL3), a common KCTD partner in protein ubiquitination/degradation (13, 23–25). While insightful structural and functional studies have provided important clues on the mechanism of action of cluster 1A members (15, 26–28), clear structure/function relationships have not yet been established for KCTD1 and its close homolog KCTD15. Several literature reports have clearly demonstrated that the close homologs KCTD1 and KCTD15 recapitulate at a smaller scale the peculiar functional diversification of the KCTD family, due to their involvement in quite different biological processes. Indeed, it has been shown that dysregulations of these proteins are linked to neurodevelopmental disorders (4, 11, 29, 30), obesity (31–33), cancer (3, 34–40), and genetic diseases (5, 6, 11, 41, 42). Notably, KCTD1 and KCTD15 mutations have been associated with scalp-ear-nipple syndrome (SENS), Aplasia cutis congenita, kidney fibrosis, and cardiac outflow tract abnormalities ((11) and references therein). We and others have recently shown that the two proteins cooperate by forming functional heterocomplexes (11). Although these disease-causing mutations are generally located in the folded regions of the KCTD1 protein (41, 43), somehow surprisingly, one of them Pro20Ser is located in the presumably unfolded N-terminal region that precedes the BTB domain (preBTB). To gain insights into the intricate role of the KCTD1 protein in different physiopathological contexts, we here present the crystal structure of the full-length disease-causing mutant P20S. The global pentameric structure of the protein presents an intricate architecture in which adjacent chains exchange the entire BTB domain to generate a closed domain swapping (44, 45). Uncommon interactions such as helix-helix discontinuous association and intermolecular PPII/PPII recognition domains characterize the interdomain contacts. The combination of these experimental data with computational studies provides a clear explanation of the puzzling pathogenicity of the P20S KCTD1 mutation. More importantly, the presence of a potassium ion bound to the central cavity of the CTD pentameric assembly provides insights into the role of KCTD1 and the other members of the family in metal homeostasis. Finally, we also describe the predicted model for a long-form of the protein which is conjugated with a Crypton transposon (46).

## 2. Results

### 2.1 Structural characterization of the disease-causing mutant P20S of KCTD1

To gain insights into the multiple biological roles of KCTD1, we undertook a structural characterization of the full-length protein (residues 1-257). All the attempts to grow crystals suitable for crystallographic investigations of the wild-type of the protein were unsuccessful. On the other hand, crystals amenable to structural studies were obtained for the full-length form of the disease-causing mutant P20S (KCTD1^P20S^). While this investigation was in progress, a structure of a truncated form (residues 28-257) of KCTD1 was reported in the Protein Data Bank (PDB) (ID 6S4L - https://doi.org/10.2210/pdb6S4L/pdb) with no publication hitherto associated. In the following paragraphs, a top-down description of the KCTD1^P20S^ structure is reported also in comparison with the previously reported 6S4L PDB structure and AlphaFold here predicted models.

#### 2.1.1 The intricate architecture of the KCTD1 pentamer: evidence for a close domain swapping

Despite the moderate resolution of the present study (2.6 Å), the electron density is well-defined for most of the residues of the protein. The disordered residues with no detectable electron densities correspond to the terminal regions (residues 1-18 and 242-257) and the loop 177-186. The inspection of the electron density also discloses structured regions at the N-terminal side of the BTB domain (preBTB), expected to be fully disordered. A well-defined density for the region 18-29 characterizes three chains, whereas the preBTB region is fully disordered in the other two chains (Fig. S1).

The global structure of the KCTD1^P20S^ pentamer is characterized by an intricate architecture in which each of its folded elements (preBTB, BTB, and CTD) establishes strong inter-rather than intra-subunitinteraction. Indeed, the individual subunits mutually exchange domains to generate a closed pentameric 3D-domain swapping motif (Fig. 1). The swapping element is the BTB domain, as this motif is exchanged by two adjacent chains. In this swapping motif, the BTB and the pre-BTB/CTD domains of one chain interact with two distinct adjacent chains (Fig. 1B).

**Figure 1.**
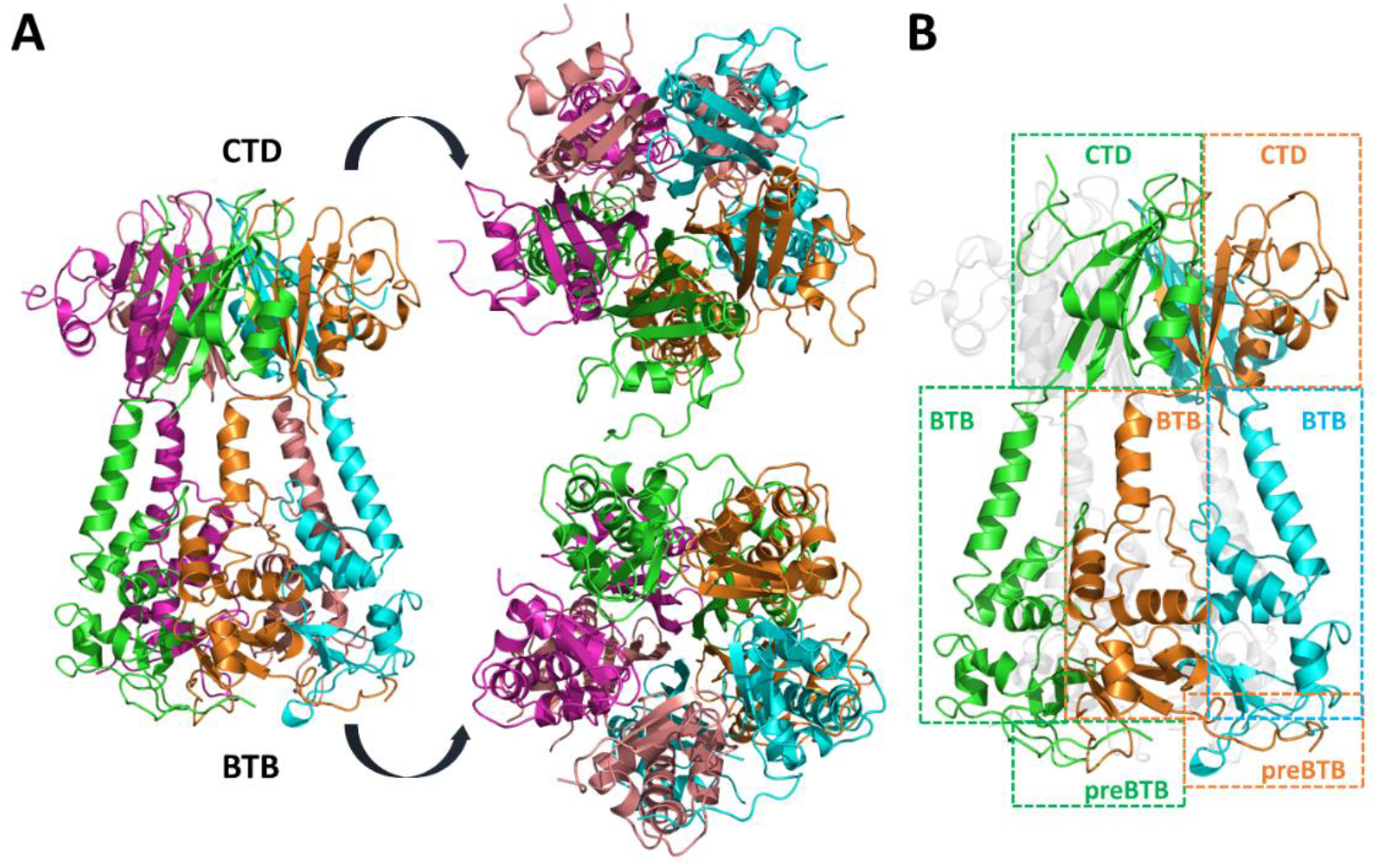
The intricate architecture of the disease-causing mutant P20S of KCTD1. Different views of the pentamer colored by chain (A) and schematic representation of its closed 3D-domain swapping motif (B).

Although 3D-domain swapping is a widespread phenomenon in protein three-dimensional structures (45, 47), the structural complexity of KCTD1 in terms of the oligomeric state and the size of the swapped element is rather unusual. A deep inspection of the interactions that stabilize the 3D-domain swapped pentamer of KCTD1^P20S^ indicated that both the preBTB/BTB and BTB/CTD interfaces rely on intriguing and somehow unusual association modes. As anticipated above, the preBTB domain, which was expected to be unfolded and was even removed in a previous crystallographic study of the isolated BTB domain of KCTD1 to facilitate the crystallization, is in part well-folded in KCTD1^P20S^. Indeed, in three out of the five chains of the pentamer, the preBTP region embedding residues 19-25 adopts a polyproline II (PPII) structure which protrudes to a hosting cavity of the BTB domain of an adjacent chain (Figs. S1 and 2).

The preBTB/BTB interface is stabilized by three hydrogen-bonding interactions that involve the main chain of Thr21 (O atom) and Ala23 (N atom) of the PPII fragment and both the O and N atoms of Ile66 on the BTB domain. The interaction is further stabilized by an H-bond formed by the Thr21 side chain with the oxygen atom of the residue Gly62. The analysis of the PDB models of the KCTD1 and its BTB domain (PDB IDs 5BXB and 6S4L) in which the pre-BTB region was truncated shows that the structure of the PPII recognition motif on the BTB of this protein is virtually identical to the one here detected.

This suggests that this BTB cavity is appropriately pre-organized to anchor the PPII helix. It is important to note that the SENS-causing Pro20Ser mutation that characterizes the present structure likely reduces the PPII propensity of the preBTB region, likely destabilizing its interaction with the BTB domain (see below the MD studies). A discontinuous helix formed by the helices 118-136 of the BTB domain and 203-213 of the CTD domain characterizes the inter-molecular BTB-CTD interface (Fig. 2). The interaction is stabilized by the peculiar juxtaposition of negative and positive poles of the dipoles of the two helices (+/-dipolar association). Indeed, the negative pole of the C-terminus of the BTB helix is associated with the positively charged N-terminus of the helix of the CTD. The BTB-CTD interface is further stabilized by interactions made by the charged residues that typically counterbalance the charge of the helix dipoles. In this scenario, strong electrostatic interactions are established by the side chains of Arg138 and Glu216 (Fig. 2). Moreover, the side chain of Arg138 interacts with the O atom of Gly214. Finally, a main chain – main chain H-bond is formed between Arg138 (N atom) and Gln212 (O atom). Similar BTB-CTD interactions are present in the truncated form of the protein deposited in the PDB (PDB ID 6S4L) (Fig. S2A).

**Figure 2.**
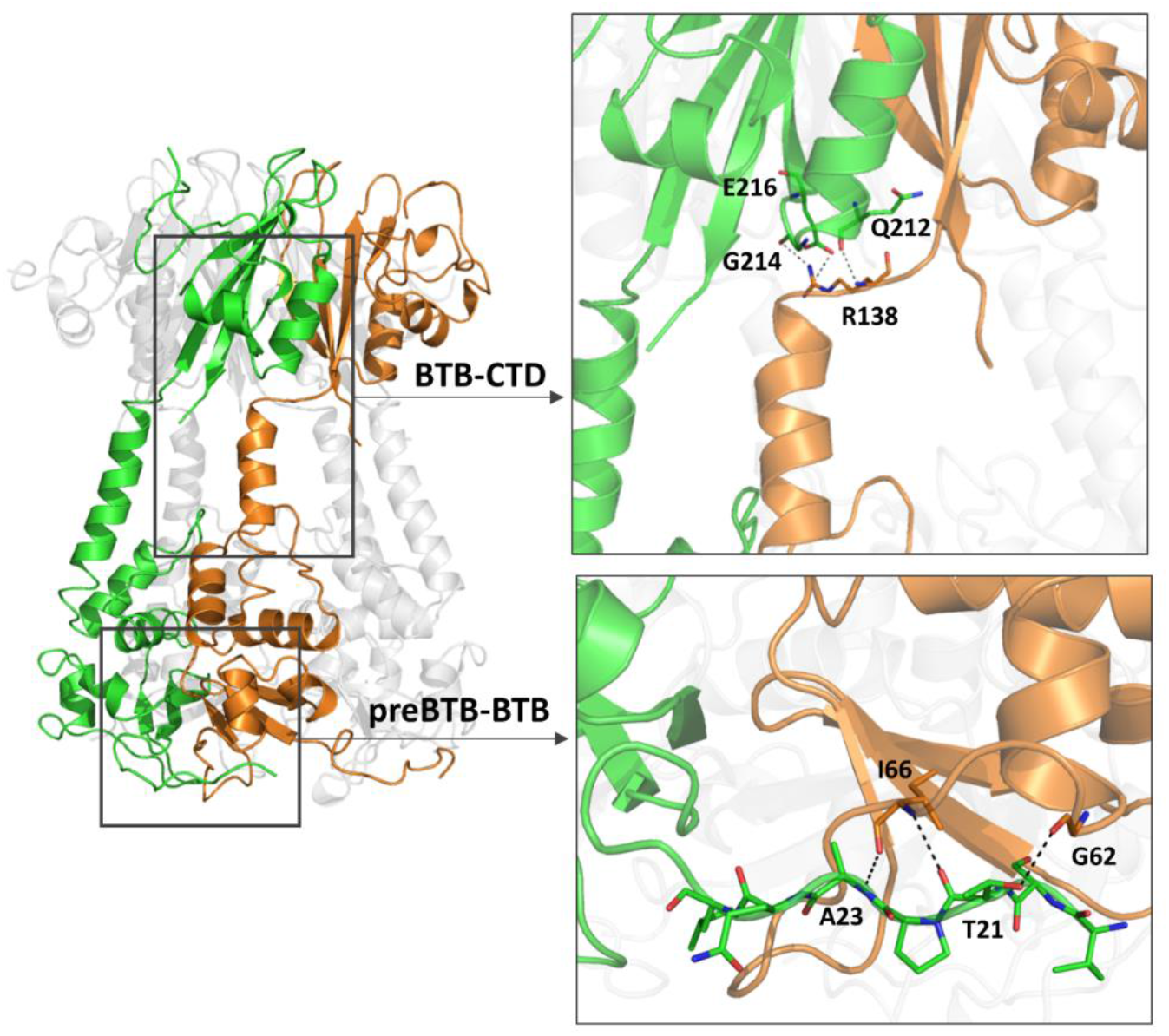
Interdomain (PreBTB-BTB and BTB-CTD) interfaces that stabilize the domain-swapped pentamer in the crystal structure of KCTD1^P20S^. Residues involved in H-bonding interactions are shown as sticks.

#### 2.1.2 The CTD domain: fold and metal binding

In line with the previous AlphaFold prediction and with the PDB structure of the truncated form of KCTD1 (PDB ID 6S4L), five copies of the CTD domain assemble to form a pentameric structure that resembles the common β-propeller fold (Fig. 3A).

**Figure 3.**
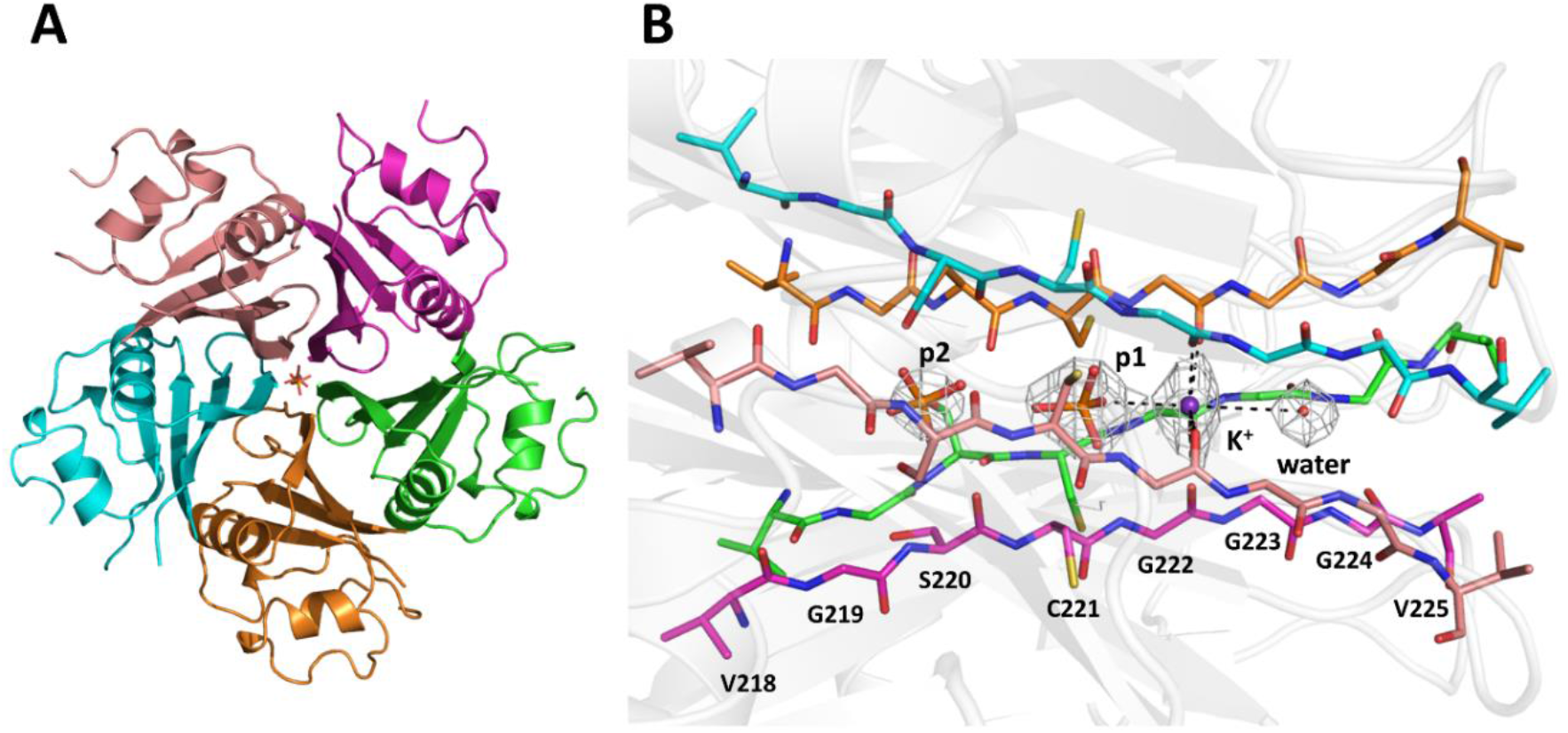
CTD domain of KCTD1^P20S^. Top view of the CTD colored by chain in the crystal structure of KCTD1^P20S^ (A). The |2Fo-Fc| electron density of the ligands (a potassium ion K^+^, a water molecule, and two phosphate groups denoted as p1 and p2) in the channel is shown as mesh at 1.3σ (B).

However, the KCTD1 CTD pentameric structure presents two main distinctive features when compared to canonical β-propellers: (a) the contribution of five distinct subunits to generate a fold that is typically formed by a single polypeptide chain, (b) the presence of two helices per subunits in addition to the β-structure, and (c) a reduced size of the central channel. Moreover, the main chain H-bond donors and acceptor of the edge β-stands, which delimitate the internal channel of the CTD pentamer, are fully exposed and not involved in H-bonding interactions. Considering the intrinsic stickiness of the exposed β-strands and the limited size of the side chains of the residues that compose the strand (219-GSCGGG-224), it is not surprising that different chemical species are bound to the internal channel formed by the assembly of KCTD CTDs. The inspection of the electron density shows the binding of two phosphate groups, a potassium ion, and a water molecule in the central channel of the pentamer (see Methods for details) (Fig. 3B). The position in the channel is dictated by H-bonding interactions with the backbone atoms of the exposed edge β-strands. In particular, the five backbone oxygen atoms of Gly222 form a plane that coordinates the potassium ion. In the channel, the average metal-oxygen distance is ∼ 2.6 Å that is in line with the expected ligand-potassium coordination distances. The coordination of the metal is completed, in the apical positions by the oxygen atom of a phosphate group (p1, K^+^-O distance of 3.2 Å) and a water molecule (K^+^-O distance of 3.8 Å). A second phosphate group (p2) directly bound to p1 closes the channel (Fig. 3B).

### 2.2 The intriguing ability of AlphaFold to predict the structure of KCTD1 and its long isoform

The determination of the crystallographic structure of KCTD1^P20S^ provides the opportunity to assess the reliability of a previously reported predicted AlphaFold (AF) model of KCTD1 and to generate models of the wild-type protein as well as of its long isoform (865 residues – UniProtKB A0A2U3U043) that also contains a tyrosine recombinase (YR) element of Crypton DNA transposons.

In a comprehensive analysis of putative three-dimensional structures of all members of the KCTD proteins, we generated the pentameric AF structure of KCTD1 regions that were expected to be folded (BTB+CTD). In line with the well-known ability of the algorithm to predict protein structures, for the protein portion embodying the BTB and the CTD domains (residues 30-239), the AF model is very similar to the here described crystallographic one (RMSD deviation computed on the C^α^ atoms of 1.21 Å). The discovery that the preBTB region of the protein is folded and contributes to the pentamer stability prompted us to carry out predictions on the full-length wild-type variant of KCTD1. As shown in Figures S2B, the generated AF model can capture, not only the BTB-CTD association mode but also the preBTB-BTB interactions. Even more surprising is the observation that the prediction of the structure of a single chain of KCTD1 provides the correct folding and orientation of the preBTB region observed in the crystal structure (Fig. S2C-D), despite the absence and an adjacent hosting BTB monomer. This finding may suggest that the BTB/preBTB orientation observed in the single chain could be a recurrent motif that the algorithm learned in the training process. However, attempts to interrogate structural databases using the preBTB region and its own BTB as the search fragment did not yield positive results. In addition to the standard KCTD1 protein, which is made of 257 residues, four other isoforms have been reported (https://www.uniprot.org/uniprotkb/Q719H9/entry) and detected at the protein level. Two of them (UniProtKB J3QLL6 and J3KSG1) correspond to smaller portions (100 and 208 residues) of the canonical forms, while another (UniProtKB J3QRK1) is similar to the main isoform as it contains and lacks a few extra residues at the N-terminus and C-terminus, respectively (260 residues). On the other hand, the fourth isoform (UniProtKB A0A2U3U043), which is much longer containing 865 residues, embodies other functional domains in addition to the preBTB, BTB, and CTD ones. Indeed, it contains a DUF3504 domain which presents some sequence similarities with Cryptons, a unique class of DNA transposons using tyrosine recombinase (46). Notably, this domain is not present in any of the KCTD15 isoforms. As no structural data is available for the DUF3504 domain, we analyzed the AF model of this long variant of KCTD1 (Fig. 4). The AF structure highlights the presence of two independent folded regions: the first (residues 167-470) including the DUF domain (residues 286-466), and the second, located at the C-terminus, corresponding to the canonical isoform. The first structural region is composed of two folded domains (residues 170-273 and 286-466), which, although not physically interacting, present a relative orientation that is assessed as reliable by AF based on the predicted aligned error map (Fig. 4B). This puzzling observation may suggest that this is a preferential orientation of similar domains present in structural databases. As shown in Figure 4C, the relative orientation of the two domains of this folded unit of the long form of KCTD1 closely resembles that found in a DNA complex of the Cre recombinase (PDB ID 5CRX) (48), which is a member of the integrase family, despite the absence of any sequence similarity between the protein. Evidently, the reliability of the orientation of the two domains in the KCTD structure that emerged from the AF self-assessment derives from the presence in the PDB of related situations. This finding demonstrates that AF predictions may provide, in addition to reliable prediction of the protein folds and of direct protein-protein interactions, clues on the structural partnerships mediated by other biomolecules.

**Figure 4.**
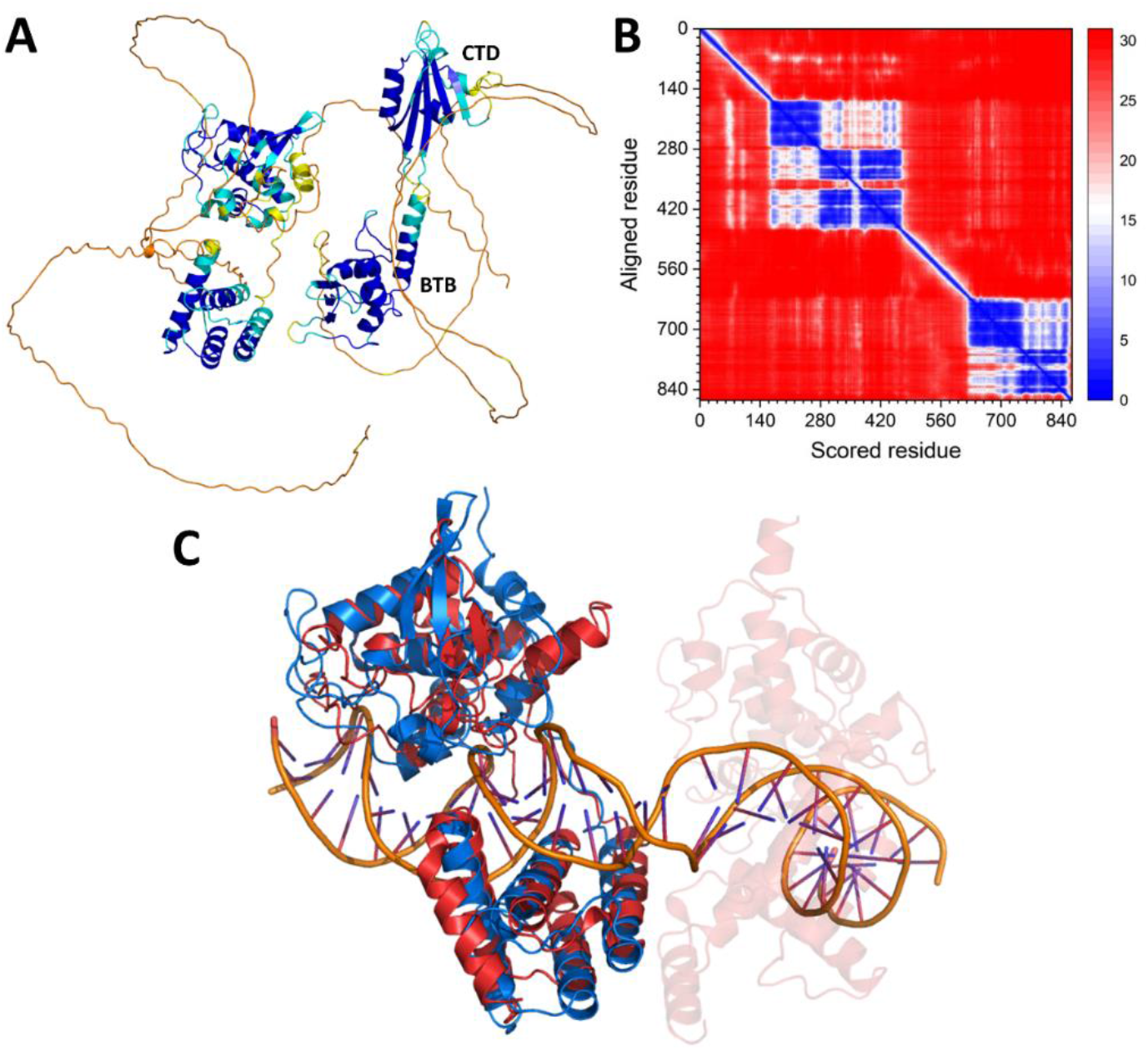
AF predicted structure of the long isoform of human KCTD1 (UniProtKB A0A2U3U043). Cartoon representation (A) and PAE matrix (B) of the KCTD1 isoform colored according to the AF per-residue confidence metric (pLDDT). Structural alignment of the N-terminal region (residues 170-466) of the KCTD1 isoform with Recombinase cre (UniProtKB P06956, PDB ID 5CRX).

### 2.3 Molecular dynamics simulations provide a dynamic view of the KCTD1 pentamer

To gain further insights into KCTD1 functions and dysregulations, we performed atomistic MD simulations on the wild-type and the Pro20Ser mutant of the protein. As detailed in the Methods and Materials section, we performed two replicas each for the wild-type and the mutant using the AF and the crystallographic structures, respectively, as the starting model.

As demonstrated by the structural indicators commonly used to monitor the trajectory of the MD simulations, all replicas achieved stable states in the first 20 ns, with RMSD values of the trajectory structures versus the starting models mostly confined in the 1.5 to 2.0 Å range (Fig. S3). The stability of the MD structures was also corroborated by the analysis of other indicators such as secondary structure and gyration radius (data not shown). A comparative analysis of the MD data for the wild-type and the mutant highlights significant differences. As shown in Fig. S4, the residue Pro20 of the wild-type KCTD1 almost uniquely adopts PPII conformations whereas Ser20 of the mutant frequently shifts between PPII and extended conformational states. This behavior has consequences for the persistence of the interactions that stabilize the preBTB-BTB association. Although, in line with the crystallographic data and the AF predictions, these interactions are generally preserved, their conservation is higher in the wild-type compared to the mutant (Fig. 5). These data indicate that the preBTB/BTB association is partially perturbed in the mutant with the likely destabilization of the pentameric assembly.

**Figure 5.**
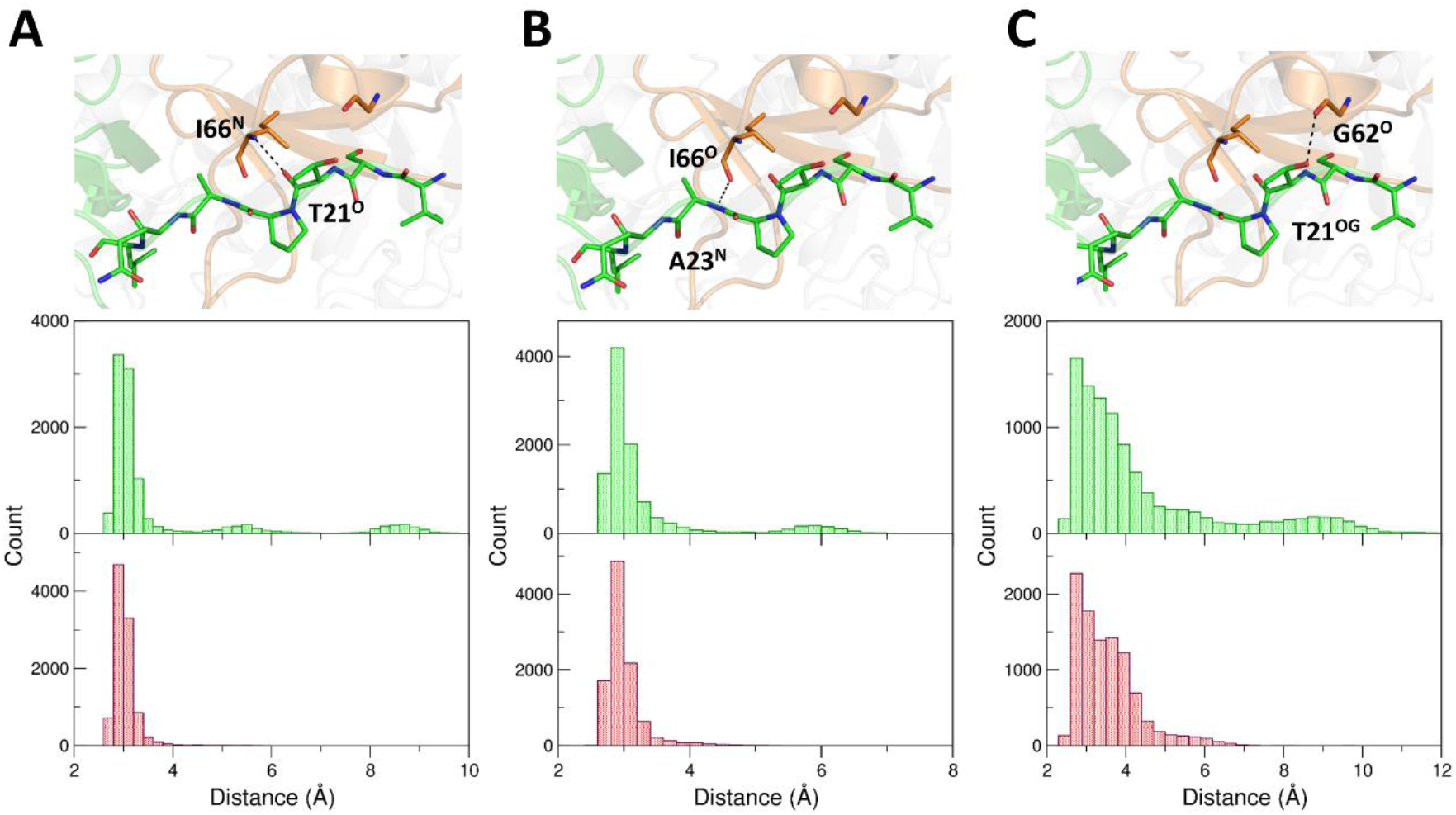
PreBTB-BTB interaction in the MD simulations. Comparative analysis of the distances that stabilize the preBTB-BTB interdomain contacts in the MD simulations of the experimental KCTD1^P20S^ structure (green) and of the predicted KCTD1 structure (red): T21O-I66N (A), A23N-I66O (B), and T21OG-G62O (C).

The availability of MD simulations, which were carried out in water without the addition of potassium or phosphate ions that could interact with the protein, also provided us with the possibility to evaluate the intrinsic dynamics of the central channel of the CTD. As shown in Figure S5, minimal variations of the channel are observed throughout the simulation, suggesting that the channel is pre-organized to anchor the ions in the positions we detected in the crystal structure.

## 3. Discussion

KCTDs represent a protein family whose members play an active role in several important physiopathological processes. Despite this, the definition of the mechanisms underlying their many functions is still limited. Here we gained detailed structural and dynamic data on one of the most important members of the family, KCTD1, by combining crystallographic analyses, prediction studies, and molecular dynamics simulations. The global architecture of the protein is characterized by an intricate association of five subunits in a pentameric structure whereby each of the three protein subunits (preBTB, BTB, and CTD) contributes to the pentamer stability by making intermolecular, rather than intramolecular, interactions. The mutual exchange of structural units among the different subunits generates an unusual pentameric closed 3D-domain swapping. This particular association of the five chains is not just a structural curiosity but holds important consequences for the mechanism of the pentamer (de)stabilization, as modifications inside a single unit are likely to affect interchain rather than intrachain interactions. This observation is suited to explain on a structural basis the effects induced by the mutation of Pro20 to Ser, which has been reported as a causative event of the SENS (43). Data here presented show that this mutation site does not belong, as previously believed, to an unfolded region but it is embedded in a PPII motif that is deeply involved in intermolecular interactions that stabilize the global pentameric assembly. As corroborated by MD simulations, the substitution of the proline weakens the interaction of the PPII region with the BTB of an adjacent chain and destabilizes the global protein structure. This consideration is in perfect agreement with the observation of a partial destabilization of the KCTD1 structure induced by this mutation or by the truncation of the N-terminal preBTB region of the protein. This finding explains the observed destabilization of the pentamer that has been associated with a higher propensity of the protein to aggregate and become unable to bind its functional partner TFAP2A (41).

The observation that the preBTB region plays an active role in the stabilization of the protein goes well beyond the Pro20Ser mutation. Indeed, as shown in Figure S6, other disease-related mutations affect the preBTB/BTB recognition sites, involving, in particular, residues Gly62 and Ile66 that play a direct role in the interaction (5, 49). Interestingly, with the same conceptual framework, the pathological phenotype detected in heterozygous mice bearing the KCTD1 mutation Ile27Asn (residue 19 in the human sequence) can be ascribed to the destabilization of this interaction (50). It is worth mentioning that a similar structural arrangement, with the preBTB playing an active role in the pentamer interactions, is predicted also for KCTD15 (Fig. S7), a close homolog that cross-acts with KCTD1 both physiologically and pathologically (6). The pathological mechanism and the dominant effect of these mutations are likely related to the propensity of destabilized KCTD1 variants to form amyloid-like aggregates both in vitro (41) and in cell systems (11). It has been recently reported that mutation in the C-terminal region of the protein (Arg241Gln and Pro243Ser) may be associated with dental anomalies (51). The authors suggested that these mutations can alter the ability of the protein to establish biological partnerships and/or undergo posttranslational modifications. Although this region does not assume a rigid structure in the crystallographic structure of the protein and its AlphaFold 2 model is not reliable (Fig. S2C), its predicted structure using the newly released AlphaFold 3 reliably suggests that this region folds in a PPII motif that is inserted in the cavity formed by the C-terminal helices of the BTB domain of adjacent chains. In this scenario, the two mutations destabilize these interactions either by eliminating the contacts formed by Arg241 or by undermining the PPII motif (Fig. S8).

In more general terms, KCTD1 functionality is extremely sensitive to amino acid replacements. Indeed, as indicated by the predictions reported in the AlphaMissense database (52), except for a limited number of residues, the vast majority of replacements are likely to induce pathological effects (Fig. S9). This is also detected for the preBTB PPII region. Notably, the percentage of amino acid replacements in KCTD1 that are predicted to be pathological by AlphaMissense is 76.5. This value is particularly high if compared to the global percentage of predicted pathological replacement in the human proteome by the same algorithm (32%) (52). This observation well agrees with the remarkable conservation of the protein during the evolution. As shown in Table S1, the mouse and human sequences are 100% identical, a rather uncommon situation detected in only 2.3% of cases in large-scale statistical comparisons (53). Remarkable similarities are observed also in comparison with more distant organisms (Table S1). The extreme susceptibility of KCTD1 to amino acid replacements is likely due to its intricate architecture and propensity to aggregate (11, 41, 54).

The CTD domain, which adopts a structure that constitutes a variation of the common β-propeller fold, is similar to that previously reported in the truncated forms of the protein. The structure of the CTD pentamer has close analogies with that exhibited by the GTP cyclohydrolase 1 feedback regulatory protein (GTPCH-GFRP) (Fig. 6A).

**Figure 6.**
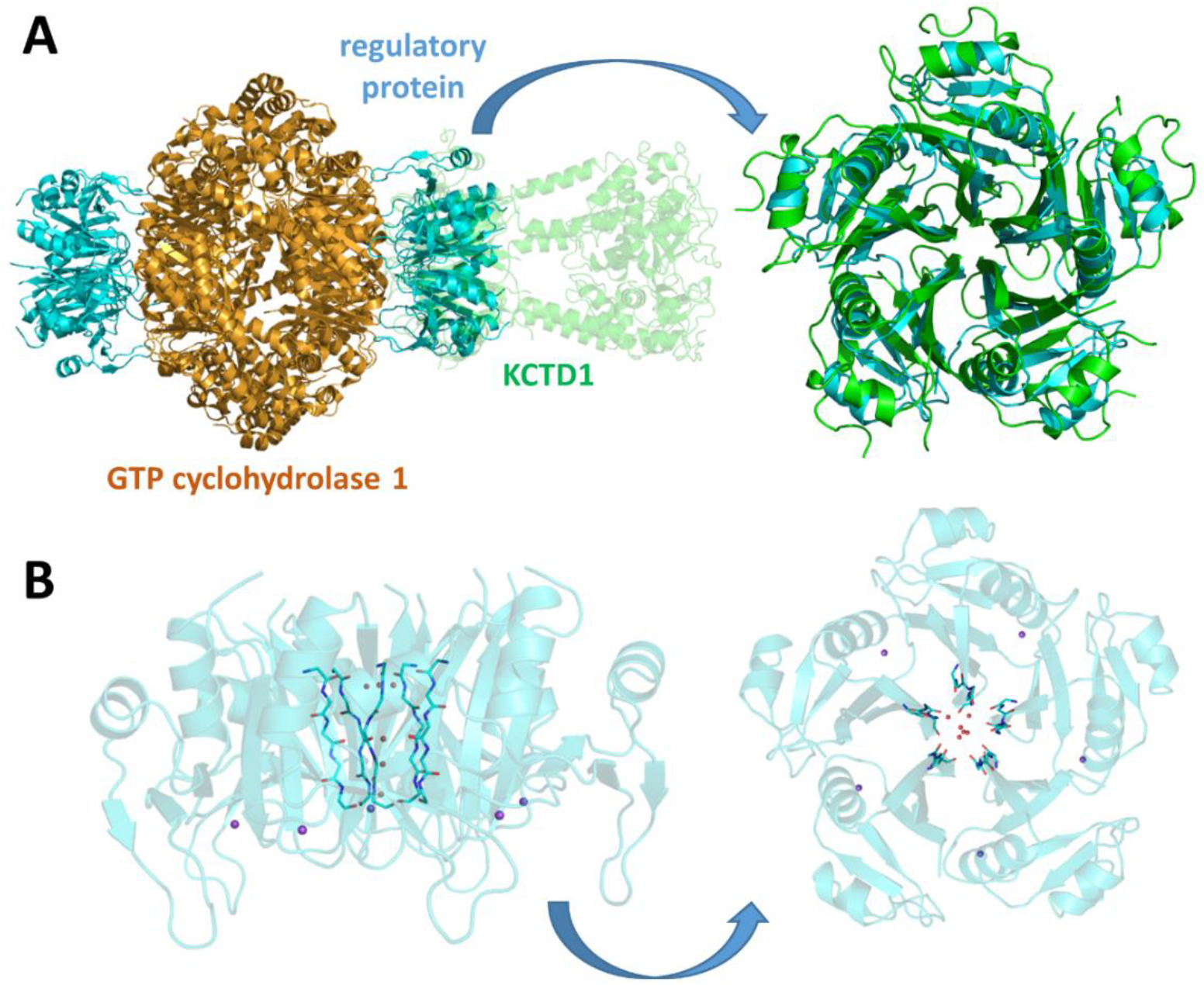
Structural analogies between KCTD1 and the GTP cyclohydrolase 1 feedback regulatory protein (GTPCH-GFRP). Structural alignment of the CTD domain of KCTD1^P20S^ with the GTP cyclohydrolase 1 feedback regulatory protein (UniProtKB P30047, PDB ID 7ALA) (A). Different views of the channel formed by GFRP. Residues forming the central channel and water molecules inside the channel are shown as sticks (only main chain atoms) and spheres, respectively. Sodium ions are shown as purple spheres.

Although global alignments of the sequence of GTPCH-GFRP with that of KCTD1 do not yield remarkable similarities, a significant sequence identity is detected for the exposed β-strand that delimitates the channel, thus indicating a common evolutionary origin of the two proteins. Despite the striking structural analogy of KCTD1 and GTPCH-GFRP, only water molecules were found in the internal channel of the regulatory protein (Fig. 6B), notwithstanding the presence of potassium ions in the crystallization medium. The fold of the KCTD CTD domain has also significant similarities with the paraflagellar rod component, a kinetoplastid-specific protein (Fig. S10). Although the paraflagellar rod component adopts a tetrameric state, its central channel is occupied by positive and negative ions, similar to KCTD1 (Fig. S10A). Moreover, the search for structural homologs of the CTD domain was conducted also considering the ensemble of all human proteins whose structure was predicted by AF using DALI (55). This survey uncovered that the protein Raftlin-2 can adopt a similar structural organization. However, in this protein, a four-blade propeller formed by a single polypeptide chain is observed (Fig. S10B). Although the biological meaning of this similarity is yet to be disclosed, it nevertheless indicates that this type of structure may be achieved using either single or multiple subunits.

As we previously anticipated using the predicted structures of KCTDs (22), the CTD domains of these proteins present significant analogies despite the marginal or absent sequence similarities between the members of different clusters. In line with these predictions, we here observe that some features of the propeller-like folding of the CTD of KCTD1 are present not only in the close homolog KCTD15 but also in distant KCTDs. In particular, the propeller-like fold with a solvent-exposed backbone of the edge strand is detected in other members of the family (see Fig. S11 for some representative examples).

In the crystallographic structure of KCTD1 here determined, the central channel, which presents a funnel-like shape, is filled with different ions. The anchoring of these ions is assured by the exposure of the oxygen and the nitrogen backbone atoms of the edge strand which is generally composed of residues with limited sizes. Due to the fivefold symmetry of the channel, planes of pentamers formed by these H-bond donors and acceptors may be identified. In KCTD1, the oxygen atom of Gly222 assures a planar penta-coordination of a potassium ion, in a manner that vaguely resembles that found in crown ethers. The coordination of the metal is completed by a phosphate group and a water molecule present in the funnel that limits its mobility. It is worth mentioning that this coordination is similar to that observed for the calcium in calmodulin (Fig. S12) and that this metal could be located in the channel with a suitable geometry without any steric hindrance. Based on these observations and considerations, the central funnel of KCTD1 seems to be specifically designed to bind the potassium and likely other metals; the other ions or water molecules bound into the channel strongly limit the mobility of the metal, thus favoring its sequestration. The ability of KCTD1 to bind metals may be related to its role in metal homeostasis.

The inspection of the structural models of KCTDs whose CTD is endowed with a propeller-like fold indicates that the funnel may have different sizes (Table S2). The data reported in the Table indicate that KCTD1/KCTD15 presents the narrowest funnels and a channel size that can directly bind metals. Rather narrow funnels are also formed by the of the 1A cluster (KCTD8/KCTD12/KCTD16). For the other KCTDs, the putative binding of the metal requires a significant reduction of the funnel.

In conclusion, considering the importance of metals in neurodevelopmental and neurologic disorders, the finding emerged in this study, that KCTD proteins are potentially able to bind metals represents an intriguing hypothesis to be investigated in further studies. Recent studies showing that the effects of the KCTD5 downregulation in mice models may be mitigated by the addition of metal chelators go in the direction here proposed.

## 4. Materials and Methods

### 4.1 Protein sample preparation

The KCTD1 protein used in this study includes residues 1-257 of the entire protein sequence (UniProtKB Q719H9) bearing the Pro20Ser mutation (KCTD1^P20S^). The corresponding encoding sequence was cloned into the pETM-11 expression vector to produce an N-terminal cleavable His-tag protein.

### 4.2 Protein expression, purification, and crystallization

The Se-Met derivative of KCTD1^P20S^ was expressed in E. coli Rosetta (DE3)2 cells in 1L of minimal media (M9) enriched with the following components: 0.4% (w/v) glucose, 1 mM MgSO4, 0.1 mM CaCl_2_, 50 µg/mL ampicillin, 33 chloramphenicol, 100 µg/mL thiamine at 37°C. After reaching an OD _600_ of 0.7, an amino acid mix (50 µg/mL of Ile, Leu, and Val and 100 µg/mL of Phe, Thr, and Lys) was added to the bacterial culture. After equilibration, 60 µg/mL of seleno-L-methionine was added and the induction was performed by adding 0.5 mM IPTG. Labeled and unlabelled KCTD1^P20S^ proteins were purified by coupling two consecutive chromatographic steps on the soluble bacterial lysate. The first step was carried out by using Ni-NTA affinity resin (Qiagen), then pooled fractions, containing KCTD1^P20S^ protein, were concentrated and loaded on the size-exclusion chromatography on S200 10/30 column equilibrated in 50 mM Tris-HCl, 150 mM NaCl, 5 % (v/v) glycerol, 0.01% (w/v) CHAPS and 2 mM DTT (pH 7.8) The homogeneity of the protein was evaluated by SDS–PAGE analysis. The molecular mass of the purified protein was checked by mass spectrometry and no proteolysis of the protein was detected. Crystallization trials were performed by using the hanging-drop method at 293 K. After a preliminary screening of the crystallization conditions using different crystallization screens (Crystal Screen I and II, Index, Hampton Research), we were able to grow good quality crystals in 1.8 M sodium chloride, 0.1 M sodium phosphate monobasic monohydrate, 0.1 M Potassium phosphate monobasic, and 0.1 M Mes monohydrate (pH 6.5) with a protein concentration of 4-6 mg/mL. Moreover, a further optimization of crystallization conditions was achieved using 0.2 M of sodium thiocyanate as an additive (Additive Formulation, Hampton Research) into the protein-precipitant drop. Good diffraction crystals were obtained for both Se-Met labeled and unlabelled KCTD1^P20S^ proteins.

### 4.3 Data collection and processing

Diffraction data for the Se-Met derivatives of KCTD1^P20S^ were collected at the ID30B synchrotron beamline at European Synchrotron Radiation Facility ESRF in Grenoble (France) at 100 K. Cryoprotection of the crystals was achieved by a fast soaking in a solution containing glycerol to a final concentration of 14% (v/v). Anomalous diffraction data were collected at the peak of absorption of the selenium (0.979 Å). The crystals belong to the space group P21 (Table 1).

**Table 1.**
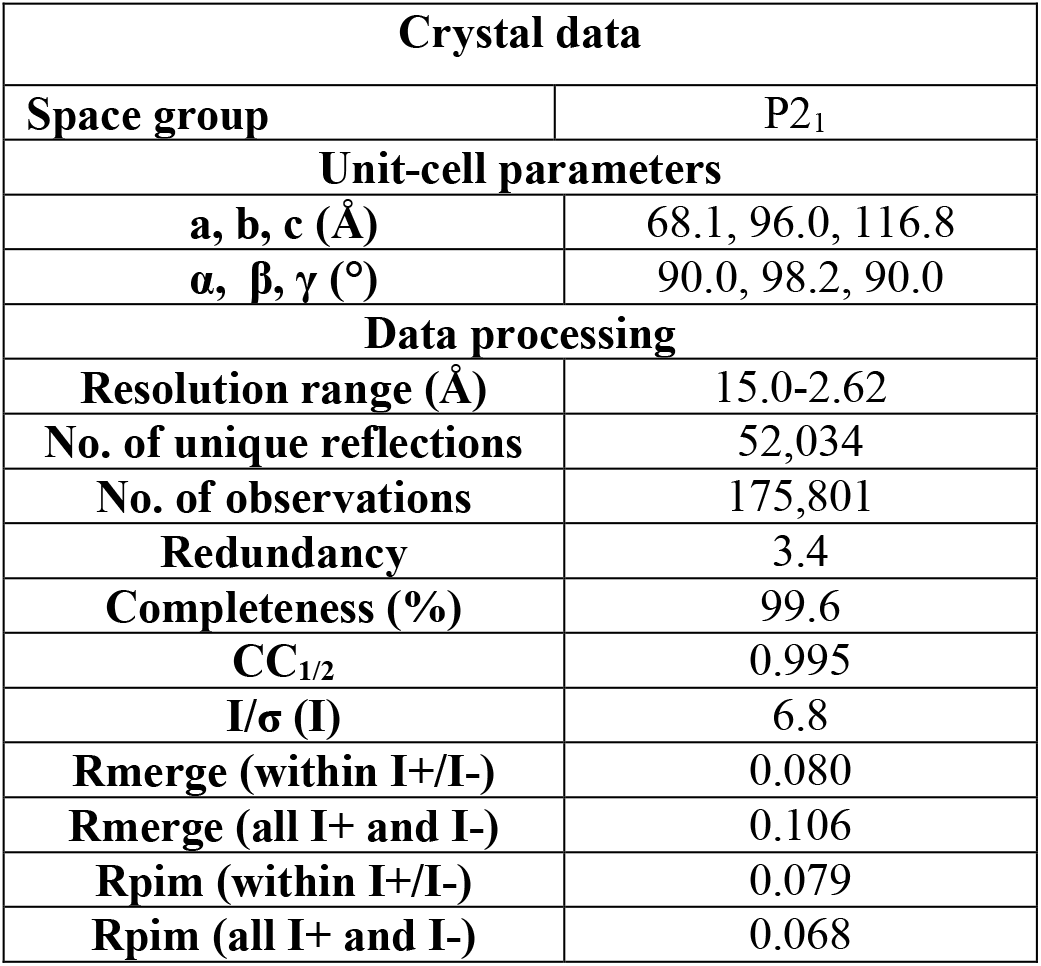
X-ray data collection statistics for KCTD1^P20S^.

### 4.4 Structure solution and refinement

The data were processed using the program XDS and the automated protocol available at ESRF (Table 1). The structure of KCTD1^P20S^ was solved using Single Anomalous Dispersion techniques and the Phenix suite (56). Crystallographic refinement of the structure of KCTD1^P20S^ was carried out against 95% of the measured data using the CCP4i program (57). The remaining 5% of the observed data, which was randomly selected, was used in R-free calculations to monitor the progress of refinement. The refinement was performed using the programs Phenix and Refmac5 (58). The refinement runs were followed by manual intervention using the molecular graphic program Coot (59), to correct minor errors in the position of the side chains. The stereochemical parameters of the refined structure (Table 2) are in close agreement with those obtained for well-refined protein structures at 2.6-Å resolution as highlighted by the program PROCHECK. In particular, 92.5% of the residues are located in the favored regions of the Ramachandran plot. The stereochemistry of the model has also been evaluated by using an innovative approach based on the monitoring of the variability of subtle details of the peptide geometry including the NC^α^C bond angle and the Δω deviation from the peptide bond planarity (60, 61). To avoid any bias, this analysis was performed on a model refined using the stereochemical libraries of Phenix not considering the conformational dependence of the geometrical parameters. The results indicate that the variability of most parameters well follows the one detected in high-resolution protein structures (Table S3 and Fig. S13). The final model was refined using the option CDL (conformation-dependent libraries) of Phenix. Atomic coordinates of KCTD1 have been deposited in the PDB with 9FQ1 identification code.

**Table 2.** Refinement statistics for KCTD1^P20S^.

| **Refinement Results** |  |
| --- | --- |
| R/R<sub>free</sub> (%) | 19.4/24.3 |
| **No. of molecules** |  |
| No. of water molecules | 82 |
| No. of residues | 1062 |
| **RMSD from ideal values** |  |
| Bond lengths (Å) | 0.006 |
| Bond angles (°) | 0.944 |
| **Average B-factors** |  |
| all atoms (Å<sup>2</sup>) | 80.0 |
| **Ramachandran plot statistics** |  |
| Favored (%) | 92.5 |
| Allowed (%) | 7.1 |
| Outliers (%) | 0 |

### 4.5 AlphaFold Predictions

Predictions of structural models were carried out using the AlphaFold v2.0 (AF) algorithm as implemented on the Colab server (https://colab.research.google.com/github/sokrypton/ColabFold/blob/main/AlphaFold2.ipynb) (62).

Predictions were carried out without considering any homologous experimental template (template_mode: none) and with three as the number of recycles and using AlphaFold-multimer v.2. The best-predicted model (rank 1) out of the five computed by AF is considered. The reliability of the AF predictions was assessed by analyzing the Local Distance Difference Test (LDDT) score and the Predicted Aligned Error (PAE) matrices reported for each predicted structure.

### 4.6 Molecular Dynamics Simulations

Fully atomistic MD simulations were performed on the KCTD1 pentamer using the GROMACS software (version 2022.3) with Amber99sb all-atom force field (63). Both the wild-type (predicted model by AF) and its P20S mutant here reported (PDB ID: 9FQ1) were considered as starting models. The protein models were solvated in triclinic boxes with TIP3P water molecules and neutralized with sodium counterions. A 10 Å cut-off was applied for Lennard–Jones interactions whereas the Particle Mesh Ewald (PME) method (0.16 nm grid spacing) was used for the electrostatic interactions (64). Bond lengths were constrained using the LINCS algorithm (65). Systems were energy minimized with the steepest descent method for 50,000 steps, and subsequently equilibrated in two phases. The temperature was raised to 300 K in 500 ps (NVT ensemble), and then the pressure was equilibrated at 1 atm in 500 ps (NpT ensemble). The Velocity Rescaling and Parrinello−Rahman algorithms were applied to control temperature and pressure, respectively. Two replica of 100 ns for each system were conducted at a constant temperature (300 K) and pressure (1 atm) with an integration time step of 2 fs. GROMACS tools and the Visual Molecular Dynamics (VMD) program (66) were used to perform structural analyses of the MD trajectories. Figures of structural models and plots were generated using the PyMOL molecular visualization program and Xmgrace v50125, respectively.

## Acknowledgments

We would like to thank Giorgio Varriale, Maurizio Amendola, Massimiliano Mazzocchi, and Luca De Luca for their technical support. We would also like to thank CINECA for computational resources (ISCRA B project KCTD-CTD ID HP10BBY7W1 and ISCRA C project AF-Koli ID HP10C52U80).

We acknowledge the European Synchrotron Radiation Facility (ESRF) for the provision of synchrotron radiation facilities and we would like to thank Andrew McCarthy for assistance and support in using beamline ID30B.

## References

1. P. J. Stogios, G. S. Downs, J. J. Jauhal, S. K. Nandra, G. G. Privé, Sequence and structural analysis of BTB domain proteins. Genome Biol 6, R82 (2005).

2. M. Skoblov, et al., Protein partners of KCTD proteins provide insights about their functional roles in cell differentiation and vertebrate development. BioEssays 35, 586–596 (2013).

3. A. Angrisani, A. Di Fiore, E. De Smaele, M. Moretti, The emerging role of the KCTD proteins in cancer. Cell Commun Signal 19, 56 (2021).

4. X. Teng, et al., KCTD : A new gene family involved in neurodevelopmental and neuropsychiatric disorders. CNS Neurosci Ther 25, 887–902 (2019).

5. A. G. Marneros, et al., Mutations in KCTD1 Cause Scalp-Ear-Nipple Syndrome. The American Journal of Human Genetics 92, 621–626 (2013).

6. K. A. Miller, et al., BTB domain mutations perturbing KCTD15 oligomerisation cause a distinctive frontonasal dysplasia syndrome. J Med Genet jmg-2023-109531 (2024). 10.1136/jmg-2023-109531.

7. K. A. Metz, et al., KCTD7 deficiency defines a distinct neurodegenerative disorder with a conserved autophagy-lysosome defect. Annals of Neurology 84, 766–780 (2018).

8. W. Wang, et al., Biallelic variants in KCTD19 associated with male factor infertility and oligoasthenoteratozoospermia. Human Reproduction 38, 1399–1411 (2023).

9. Y. Wang, H. Wang, C. Wang, Lysosomal dysfunction, autophagic defects, and CLN5 accumulation underlie the pathogenesis of KCTD7-mutated neuronal ceroid lipofuscinoses. Autophagy 19, 1876–1878 (2023).

10. M. Todisco, et al., KCTD17-related myoclonus-dystonia syndrome: clinical and electrophysiological findings of a patient with atypical late onset. Parkinsonism & Related Disorders 78, 129–133 (2020).

11. J. R. Raymundo, et al., KCTD1/KCTD15 complexes control ectodermal and neural crest cell functions, and their impairment causes aplasia cutis. Journal of Clinical Investigation 134, e174138 (2024).

12. D. M. Pinkas, et al., Structural complexity in the KCTD family of Cullin3-dependent E3 ubiquitin ligases. Biochemical Journal 474, 3747–3761 (2017).

13. A. X. Ji, et al., Structural Insights into KCTD Protein Assembly and Cullin3 Recognition. Journal of Molecular Biology 428, 92–107 (2016).

14. I. S. Dementieva, et al., Pentameric Assembly of Potassium Channel Tetramerization Domain-Containing Protein 5. Journal of Molecular Biology 387, 175–191 (2009).

15. S. Zheng, N. Abreu, J. Levitz, A. C. Kruse, Structural basis for KCTD-mediated rapid desensitization of GABAB signalling. Nature 567, 127–131 (2019).

16. W. Jiang, W. Wang, Y. Kong, S. Zheng, Structural basis for the ubiquitination of G protein βγ subunits by KCTD5/Cullin3 E3 ligase. Sci. Adv. 9, eadg8369 (2023).

17. H. Zuo, et al., Structural basis for auxiliary subunit KCTD16 regulation of the GABA B receptor. Proc. Natl. Acad. Sci. U.S.A. 116, 8370–8379 (2019).

18. D. M. Nguyen, et al., Structure and dynamics of a pentameric KCTD5/CUL3/Gβγ E3 ubiquitin ligase complex. Proc. Natl. Acad. Sci. U.S.A. 121, e2315018121 (2024).

19. D. M. Nguyen, et al., “Structure and dynamics of a pentameric KCTD5/Cullin3/Gβγ E3 ubiquitin ligase complex” (Biochemistry, 2023).

20. V. Sereikaite, et al., Targeting the γ-Aminobutyric Acid Type B (GABA B) Receptor Complex: Development of Inhibitors Targeting the K + Channel Tetramerization Domain (KCTD) Containing Proteins/GABA B Receptor Protein–Protein Interaction. J. Med. Chem. 62, 8819–8830 (2019).

21. L. Esposito, et al., AlphaFold-Predicted Structures of KCTD Proteins Unravel Previously Undetected Relationships among the Members of the Family. Biomolecules 11, 1862 (2021).

22. L. Esposito, N. Balasco, L. Vitagliano, Alphafold Predictions Provide Insights into the Structural Features of the Functional Oligomers of All Members of the KCTD Family. IJMS 23, 13346 (2022).

23. N. Balasco, L. Esposito, G. Smaldone, M. Salvatore, L. Vitagliano, A Comprehensive Analysis of the Structural Recognition between KCTD Proteins and Cullin 3. IJMS 25, 1881 (2024).

24. G. Smaldone, et al., Cullin 3 Recognition Is Not a Universal Property among KCTD Proteins. PLoS ONE 10, e0126808 (2015).

25. N. Balasco, et al., Molecular recognition of Cullin3 by KCTDs: Insights from experimental and computational investigations. Biochimica et Biophysica Acta (BBA) - Proteins and Proteomics 1844, 1289–1298 (2014).

26. J. Schwenk, et al., Native GABAB receptors are heteromultimers with a family of auxiliary subunits. Nature 465, 231–235 (2010).

27. T. Fritzius, et al., KCTD Hetero-oligomers Confer Unique Kinetic Properties on Hippocampal GABA B Receptor-Induced K + Currents. J. Neurosci. 37, 1162–1175 (2017).

28. D. C. Sloan, C. E. Cryan, B. S. Muntean, Multiple potassium channel tetramerization domain (KCTD) family members interact with Gβγ, with effects on cAMP signaling. Journal of Biological Chemistry 299, 102924 (2023).

29. S. Dutta, I. B. Dawid, Kctd15 inhibits neural crest formation by attenuating Wnt/β-catenin signaling output. Development 137, 3013–3018 (2010).

30. V. E. Zarelli, I. B. Dawid, Inhibition of neural crest formation by Kctd15 involves regulation of transcription factor AP-2. Proc. Natl. Acad. Sci. U.S.A. 110, 2870–2875 (2013).

31. L. Pirone, et al., KCTD1: A novel modulator of adipogenesis through the interaction with the transcription factor AP2α. Biochimica et Biophysica Acta (BBA) - Molecular and Cell Biology of Lipids 1864, 158514 (2019).

32. the GIANT Consortium, Six new loci associated with body mass index highlight a neuronal influence on body weight regulation. Nat Genet 41, 25–34 (2009).

33. G. Smaldone, et al., The essential player in adipogenesis GRP78 is a novel KCTD15 interactor. International Journal of Biological Macromolecules 115, 469–475 (2018).

34. A. Di Fiore, et al., KCTD1 is a new modulator of the KCASH family of Hedgehog suppressors. Neoplasia 43, 100926 (2023).

35. L. Buono, et al., A Comprehensive Analysis of the Expression Profiles of KCTD Proteins in Acute Lymphoblastic Leukemia: Evidence of Selective Expression of KCTD1 in T-ALL. JCM 12, 3669 (2023).

36. G. Smaldone, et al., KCTD15 deregulation is associated with alterations of the NF-κB signaling in both pathological and physiological model systems. Sci Rep 11, 18237 (2021).

37. G. Smaldone, et al., KCTD15 Protein Expression in Peripheral Blood and Acute Myeloid Leukemia. Diagnostics 10, 371 (2020).

38. G. Smaldone, et al., KCTD15 is overexpressed in human childhood B-cell acute lymphoid leukemia. Sci Rep 9, 20108 (2019).

39. G. Smaldone, et al., The Oncosuppressive Properties of KCTD1: Its Role in Cell Growth and Mobility. Biology 12, 481 (2023).

40. L. Coppola, et al., KCTD15 Is Overexpressed in her2+ Positive Breast Cancer Patients and Its Silencing Attenuates Proliferation in SKBR3 CELL LINE. Diagnostics 12, 591 (2022).

41. G. Smaldone, et al., Molecular basis of the scalp-ear-nipple syndrome unraveled by the characterization of disease-causing KCTD1 mutants. Sci Rep 9, 10519 (2019).

42. L. Hu, et al., KCTD1 mutants in scalp-ear-nipple syndrome and AP-2α P59A in Char syndrome reciprocally abrogate their interactions, but can regulate Wnt/β-catenin signaling. Mol Med Rep (2020). 10.3892/mmr.2020.11457.

43. A. G. Marneros, et al., Mutations in KCTD1 Cause Scalp-Ear-Nipple Syndrome. The American Journal of Human Genetics 92, 621–626 (2013).

44. L. Mazzarella, L. Vitagliano, A. Zagari, Swapping structural determinants of ribonucleases: an energetic analysis of the hinge peptide 16-22. Proc. Natl. Acad. Sci. U.S.A. 92, 3799–3803 (1995).

45. M. J. Bennett, S. Choe, D. Eisenberg, Domain swapping: entangling alliances between proteins. Proc. Natl. Acad. Sci. U.S.A. 91, 3127–3131 (1994).

46. K. K. Kojima, J. Jurka, Crypton transposons: identification of new diverse families and ancient domestication events. Mobile DNA 2, 12 (2011).

47. F. Rousseau, J. Schymkowitz, L. S. Itzhaki, “Implications of 3D Domain Swapping for Protein Folding, Misfolding and Function” in Protein Dimerization and Oligomerization in Biology, Advances in Experimental Medicine and Biology., J. M. Matthews, Ed. (Springer New York, 2012), pp. 137–152.

48. F. Guo, D. N. Gopaul, G. D. Van Duyne, Asymmetric DNA bending in the Cre-loxP site-specific recombination synapse. Proc. Natl. Acad. Sci. U.S.A. 96, 7143–7148 (1999).

49. S. Su, R. Xie, X. Ding, Y. Lin, Three Cases of Bilateral Breast Absence Associated with Familial Congenital Ectodermal Defects. CCID Volume 14, 377–383 (2021).

50. S. Kumar, et al., Standardized, systemic phenotypic analysis reveals kidney dysfunction as main alteration of Kctd1 I27N mutant mice. J Biomed Sci 24, 57 (2017).

51. C. Ruangchan, et al., Genetic Variants in KCTD1 Are Associated with Isolated Dental Anomalies. IJMS 25, 5179 (2024).

52. J. Cheng, et al., Accurate proteome-wide missense variant effect prediction with AlphaMissense. Science 381, eadg7492 (2023).

53. W. Makałowski, J. Zhang, M. S. Boguski, Comparative analysis of 1196 orthologous mouse and human full-length mRNA and protein sequences. Genome Res. 6, 846–857 (1996).

54. Z. Liu, et al., Bivalent Copper Ions Promote Fibrillar Aggregation of KCTD1 and Induce Cytotoxicity. Sci Rep 6, 32658 (2016).

55. L. Holm, DALI and the persistence of protein shape. Protein Science 29, 128–140 (2020).

56. P. D. Adams, et al., PHENIX : a comprehensive Python-based system for macromolecular structure solution. Acta Crystallogr D Biol Crystallogr 66, 213–221 (2010).

57. E. Potterton, P. Briggs, M. Turkenburg, E. Dodson, A graphical user interface to the CCP 4 program suite. Acta Crystallogr D Biol Crystallogr 59, 1131–1137 (2003).

58. A. A. Vagin, et al., REFMAC 5 dictionary: organization of prior chemical knowledge and guidelines for its use. Acta Crystallogr D Biol Crystallogr 60, 2184–2195 (2004).

59. P. Emsley, B. Lohkamp, W. G. Scott, K. Cowtan, Features and development of Coot. Acta Crystallogr D Biol Crystallogr 66, 486–501 (2010).

60. N. Balasco, L. Esposito, L. Vitagliano, Factors affecting the amplitude of the τ angle in proteins: a revisitation. Acta Crystallogr D Struct Biol 73, 618–625 (2017).

61. N. Balasco, L. Esposito, A. S. Thind, M. R. Guarracino, L. Vitagliano, Dissection of Factors Affecting the Variability of the Peptide Bond Geometry and Planarity. BioMed Research International 2017, 1–9 (2017).

62. M. Mirdita, et al., ColabFold: making protein folding accessible to all. Nat Methods 19, 679–682 (2022).

63. D. Van Der Spoel, et al., GROMACS: Fast, flexible, and free. J Comput Chem 26, 1701–1718 (2005).

64. T. Darden, D. York, L. Pedersen, Particle mesh Ewald: An N ⋅log(N) method for Ewald sums in large systems. The Journal of Chemical Physics 98, 10089–10092 (1993).

65. B. Hess, H. Bekker, H. J. C. Berendsen, J. G. E. M. Fraaije, LINCS: A linear constraint solver for molecular simulations. J. Comput. Chem. 18, 1463–1472 (1997).

66. W. Humphrey, A. Dalke, K. Schulten, VMD: Visual molecular dynamics. Journal of Molecular Graphics 14, 33–38 (1996).

