## Supplementary figures and tables for "Structural studies of KCTD1 and its disease-causing mutant P20S provide insights into the protein function and misfunction"

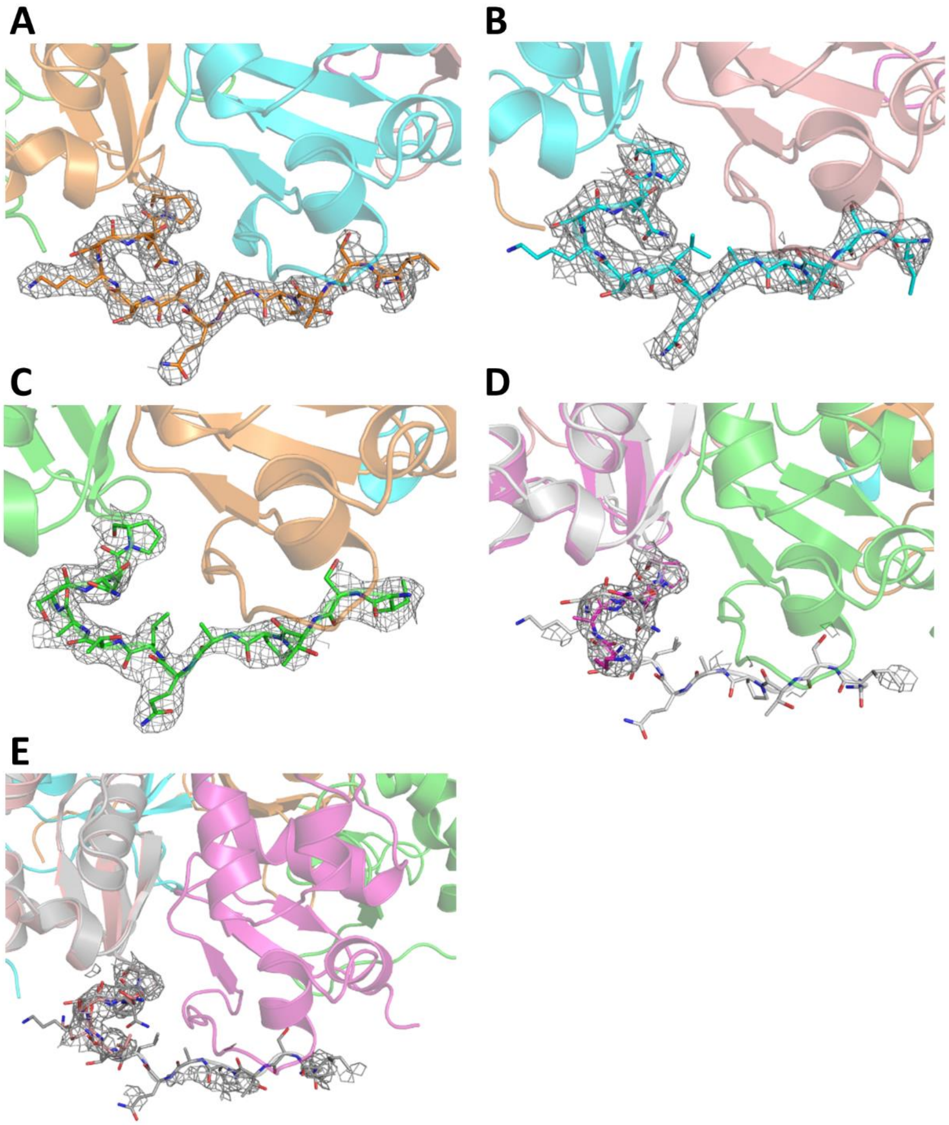


**Fig. S1.** PreBTB region in the crystal structure of KCTD1$^{P20S}$. |2Fo-Fc| electron-density maps (contoured at 1.0 σ) of the preBTB (residues 18-29) at the N-terminus of KCTD1$^{P20S}$ chains. Residues 18-29, shown as sticks, present a well-defined density in chains A (A), B (B), and C (C) whereas they are disordered in chains D (D) and E (E). In panels D and E, the absence of an interpretable electron density is highlighted by displaying the conformation that the preBTB region adopts in the chains in which it is structured (grey).

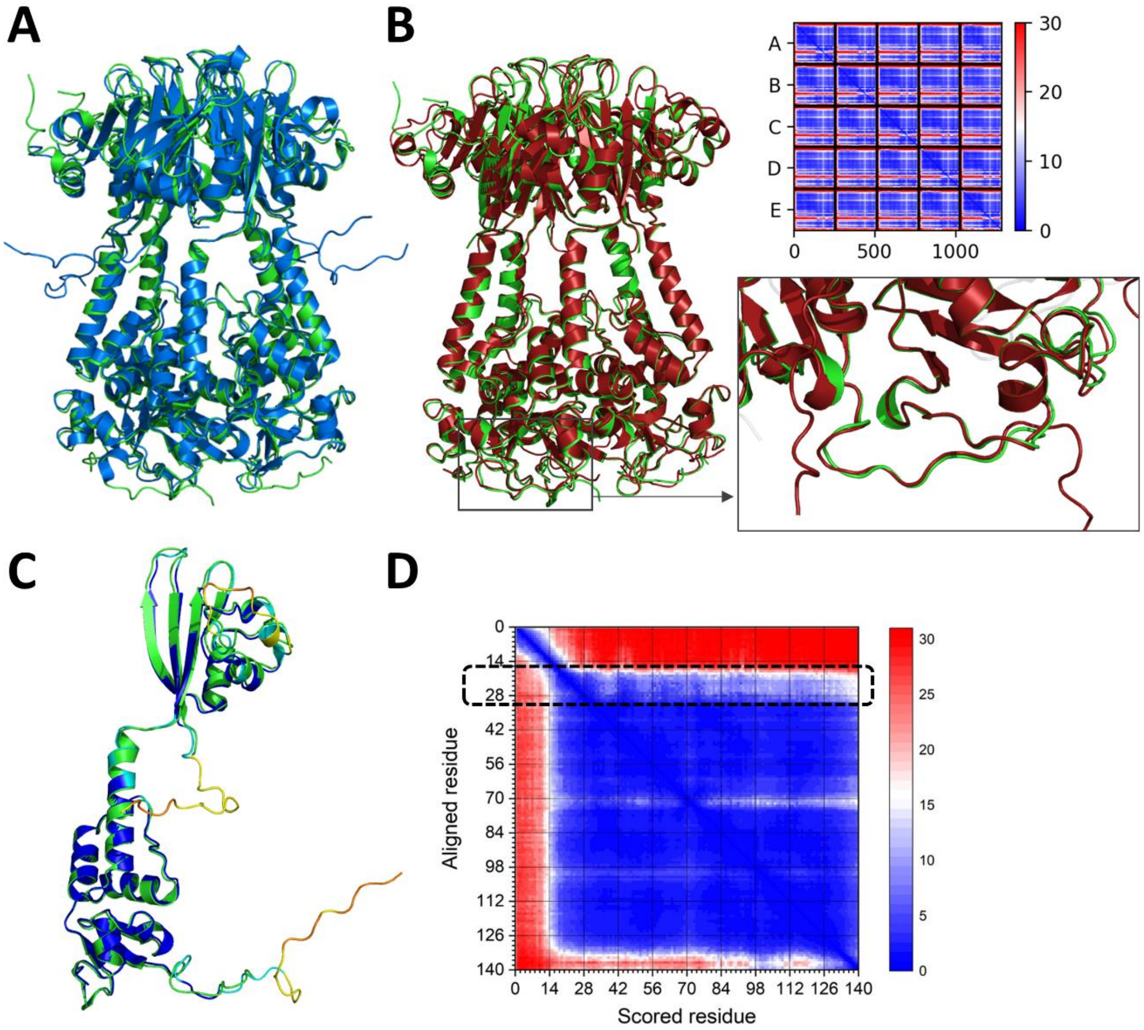


**Fig. S2.** Structural superimposition of KCTD1$^{P20S}$ pentamer (green) with the crystallographic structure (blue, PDB ID: 6S4L) (A) and the predicted AF model (red) (B) of wild-type KCTD1 pentamer. Enlargement of the BTB-PreBTB association in KCTD1$^{P20S}$ (green) and KCTD1 AF model (red). Superimposition of a single chain of KCTD1$^{P20S}$ (green) with the AF predicted structure of KCTD1 monomer (colored by pLDDT, UniProtKB Q719H9) (C). PAE matrix of the predicted AF model of wild-type KCTD1 monomer (D).

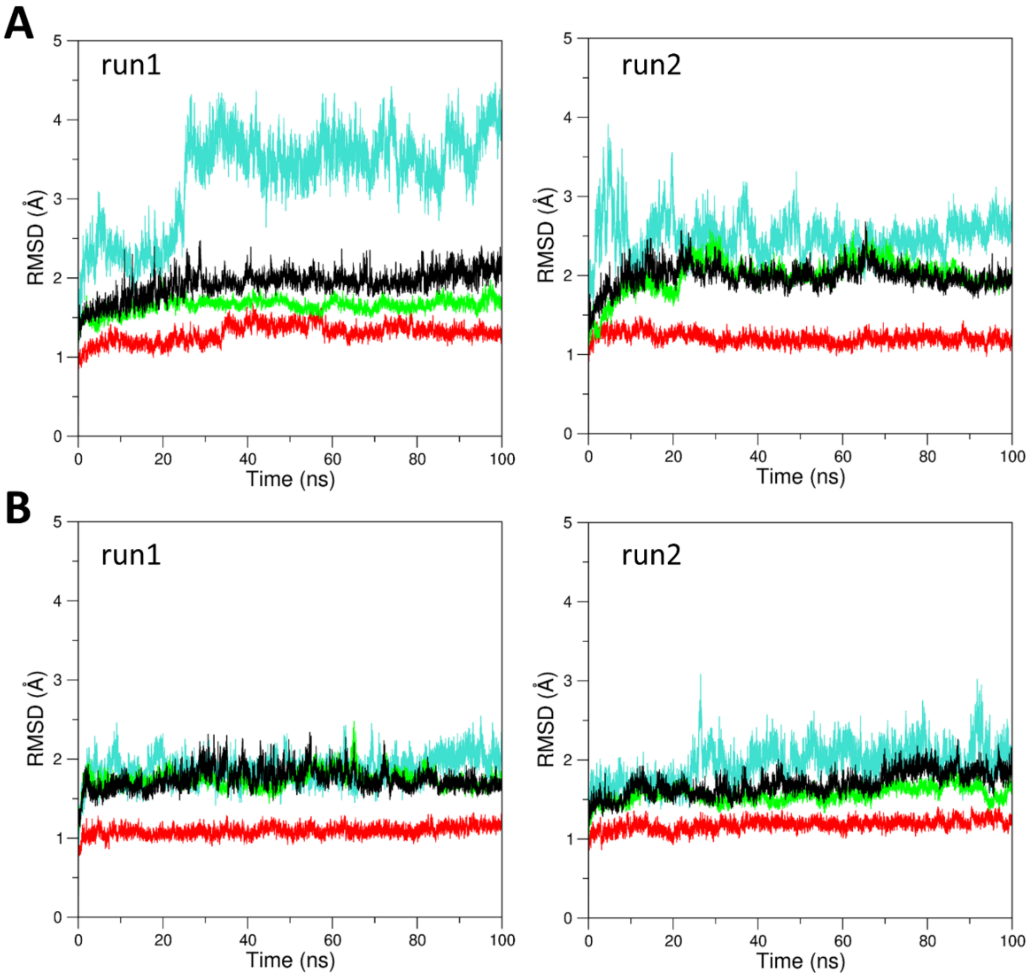


**Fig. S3.** RMSD values of trajectory structures against the starting model in the MD simulations of KCTD1$^{P20S}$ (A) and KCTD1 wild-type (B). RMSD values have been calculated on all (black), preBTB (cyan), BTB (red), and CTD (green) C$^{\alpha}$ atoms.

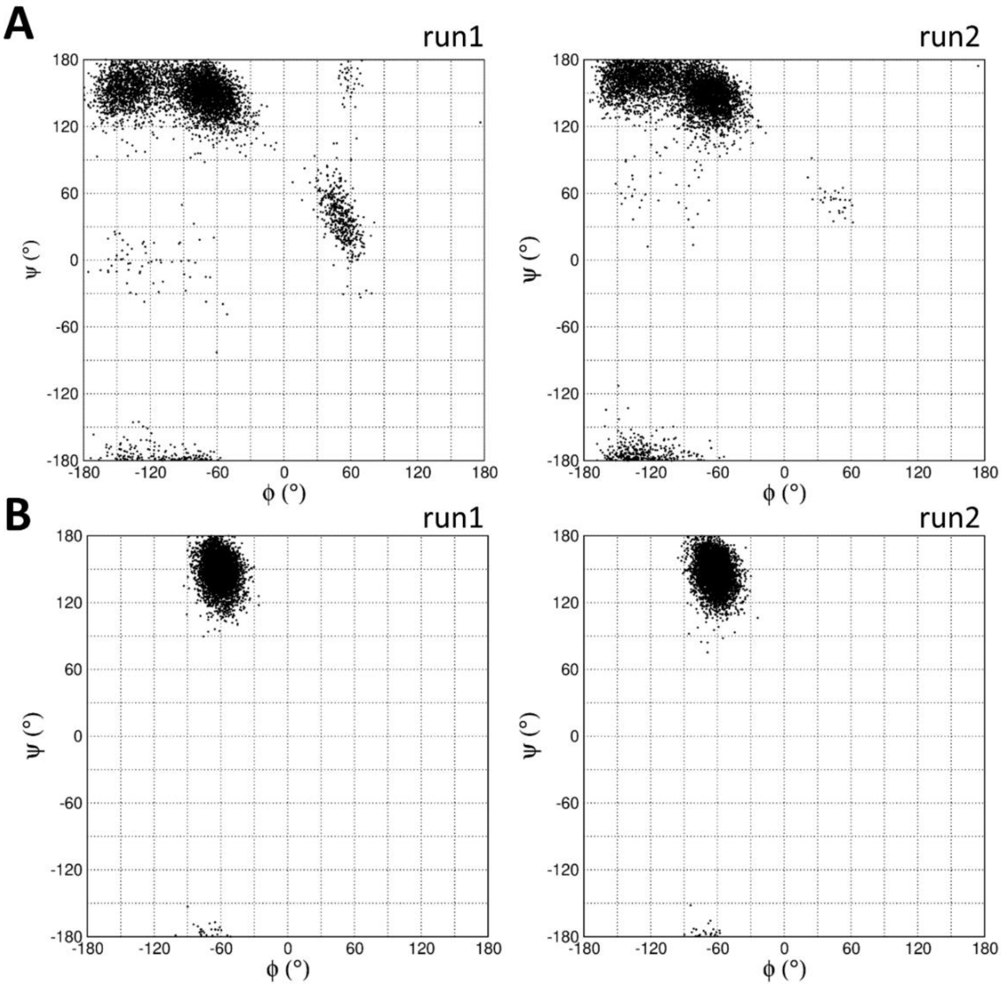


**Fig. S4.** Ramachandra plots of residue Ser20/Pro20 in the MD simulations of KCTD1$^{P20S}$ (A) and KCTD1 wild-type (B).

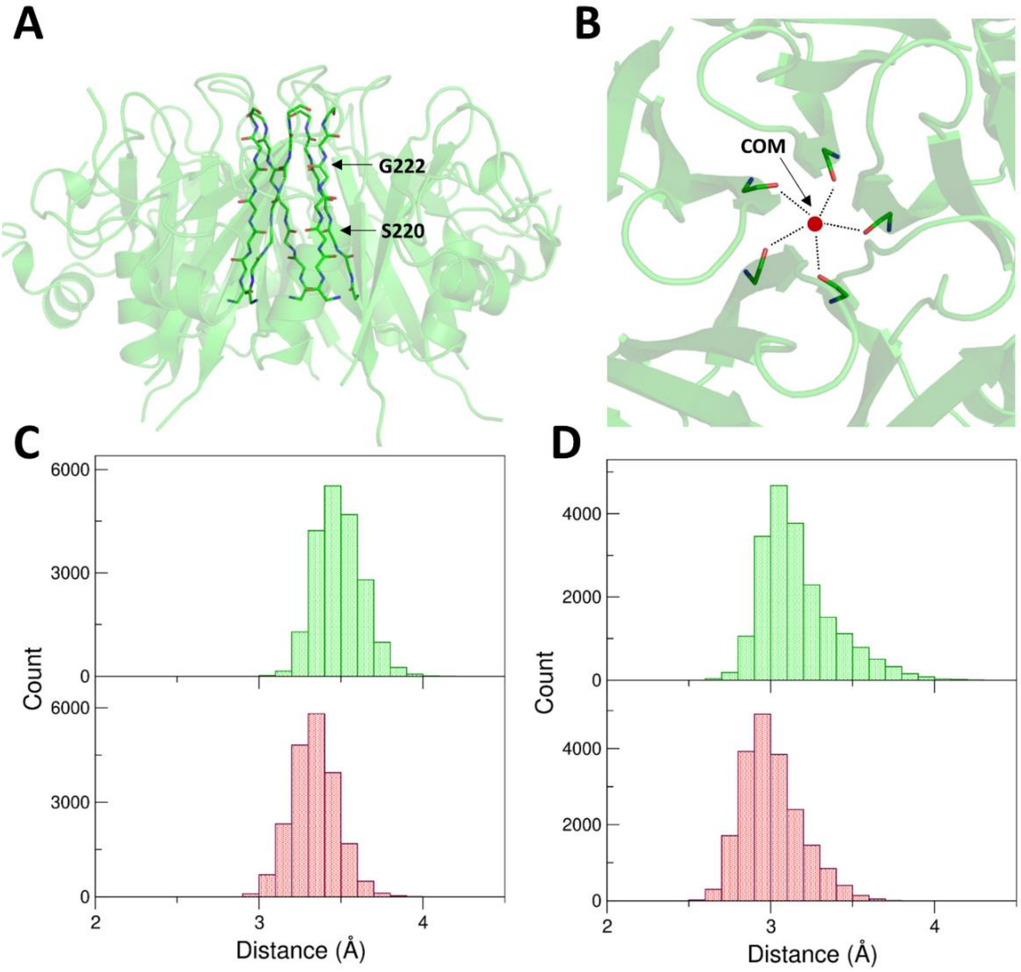


**Fig. S5.** Dynamics of the planes of the channel formed by the CTD domain. Residues that form the channel are shown as sticks (A). Schematic representation of the distances between the carbonyl oxygen atoms and the center-of-mass of the five oxygens that form the plane (B). Distribution of these distances for Ser220 (C) and Gly222 (D) residues in the MD simulations of KCTD1$^{P20S}$ (green) and KCTD1 wild-type (red).

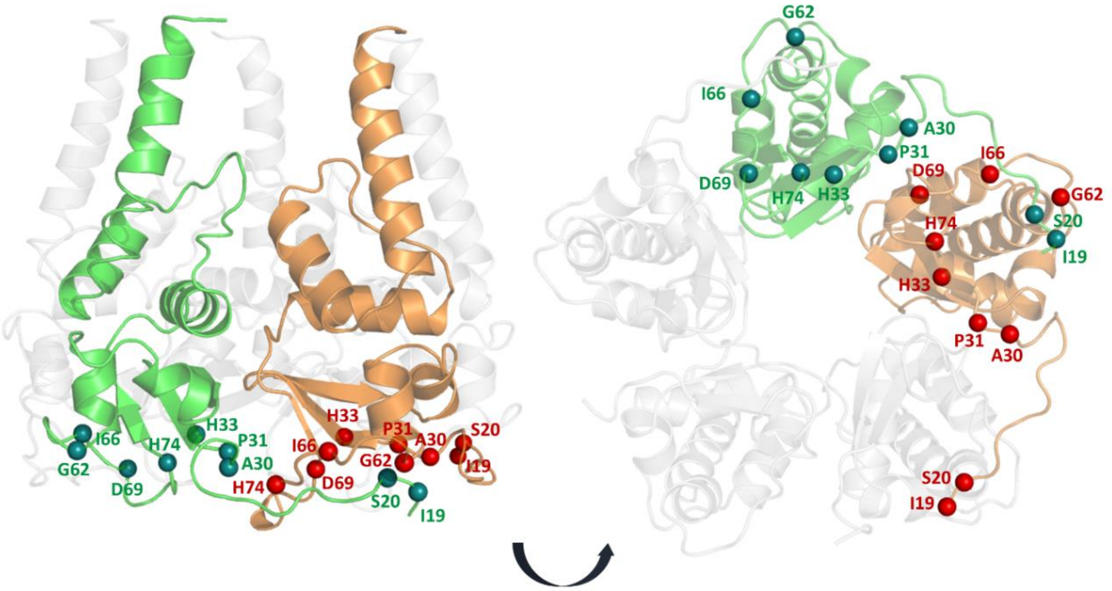


**Fig. S6.** Localization in the KCTD1 structure of disease-related amino acid mutations (residues Ile19, Ser20, Ala30, Pro31, His33, Gly62, Ile66, Asp69, and His74).

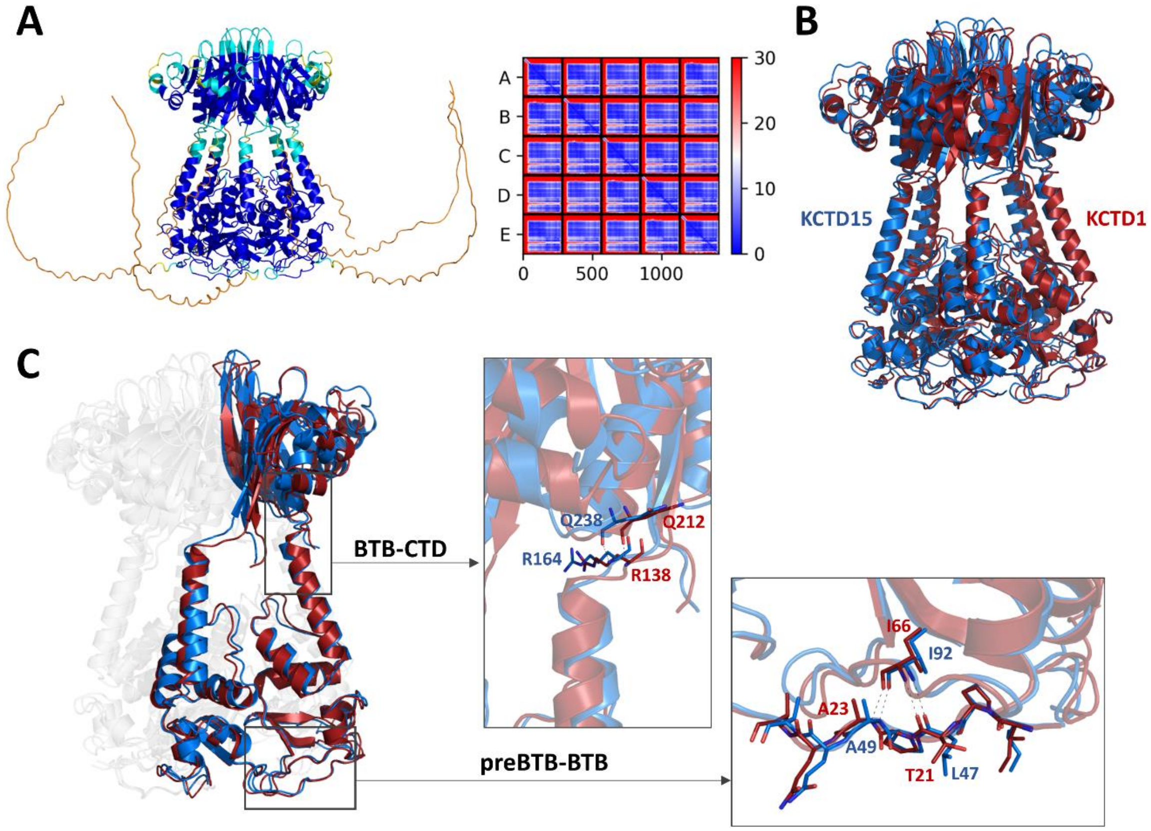


**Fig. S7.** Cartoon representation of the AF predicted model of the KCTD15 pentamer and PAE matrix (A). Structural superimposition of the AF models of KCTD1 (red) and KCTD15 (blue) pentamers (B). PreBTB-BTB and BTB-CTD interactions in KCTD1 (red) and KCTD15 (blue) AF structures (C). PreBTB residues and those involved in H-bonding contacts are shown as sticks.

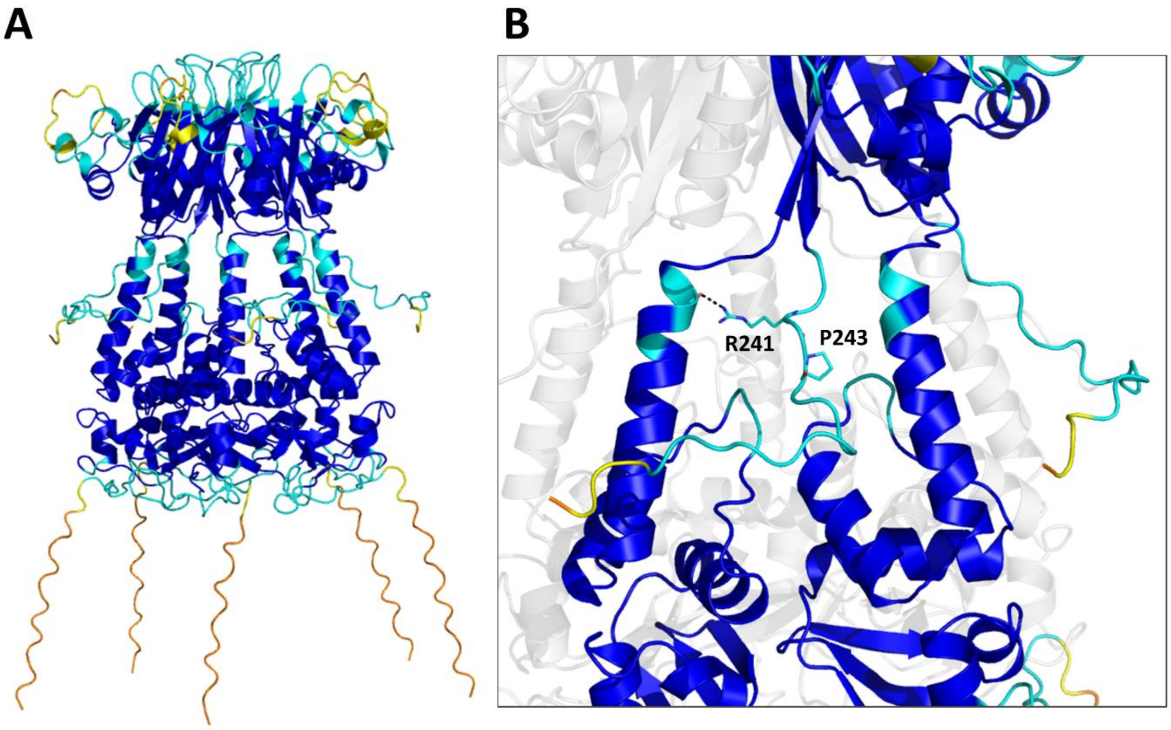


**Fig. S8.** Cartoon representation of the AF3 predicted model of KCTD1 pentamer colored following AF per-residue confidence metric (pLDDT) (A). Residues Arg241 and Pro243 in the C-terminal region are shown as sticks. The H-bonding interaction formed by Arg241 sidechain (NH1 atom) with Gly134 main chain (O atom) is shown.

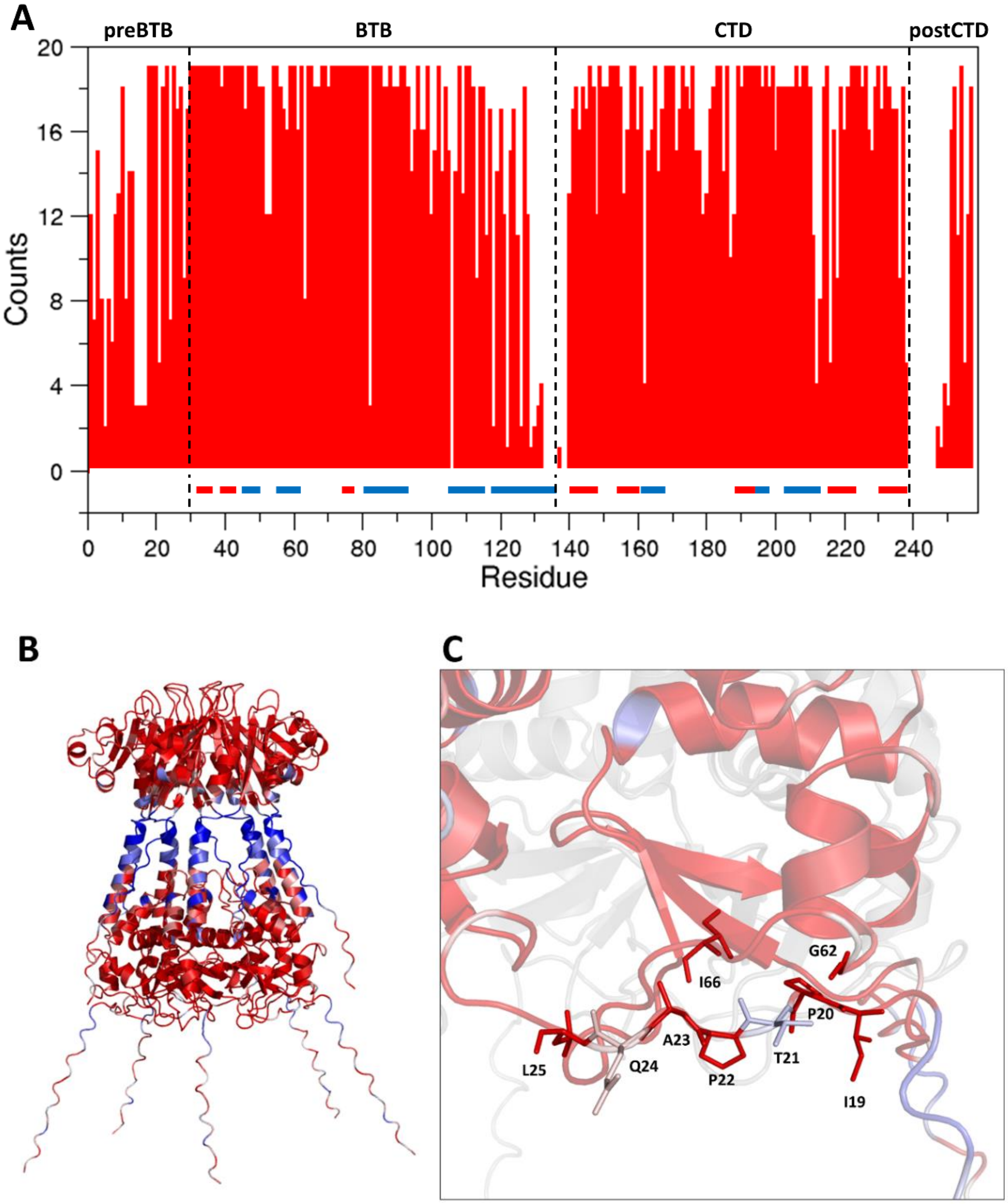


**Fig. S9.** Number of amino acid replacements in KCTD1 that are predicted to be pathological by AlphaMissense (A). Cartoon representation of KCTD1 pentamer colored from benign (blue) to pathological (red) mutations (B). Residues at the preBTB-BTB interface are shown as sticks (C).

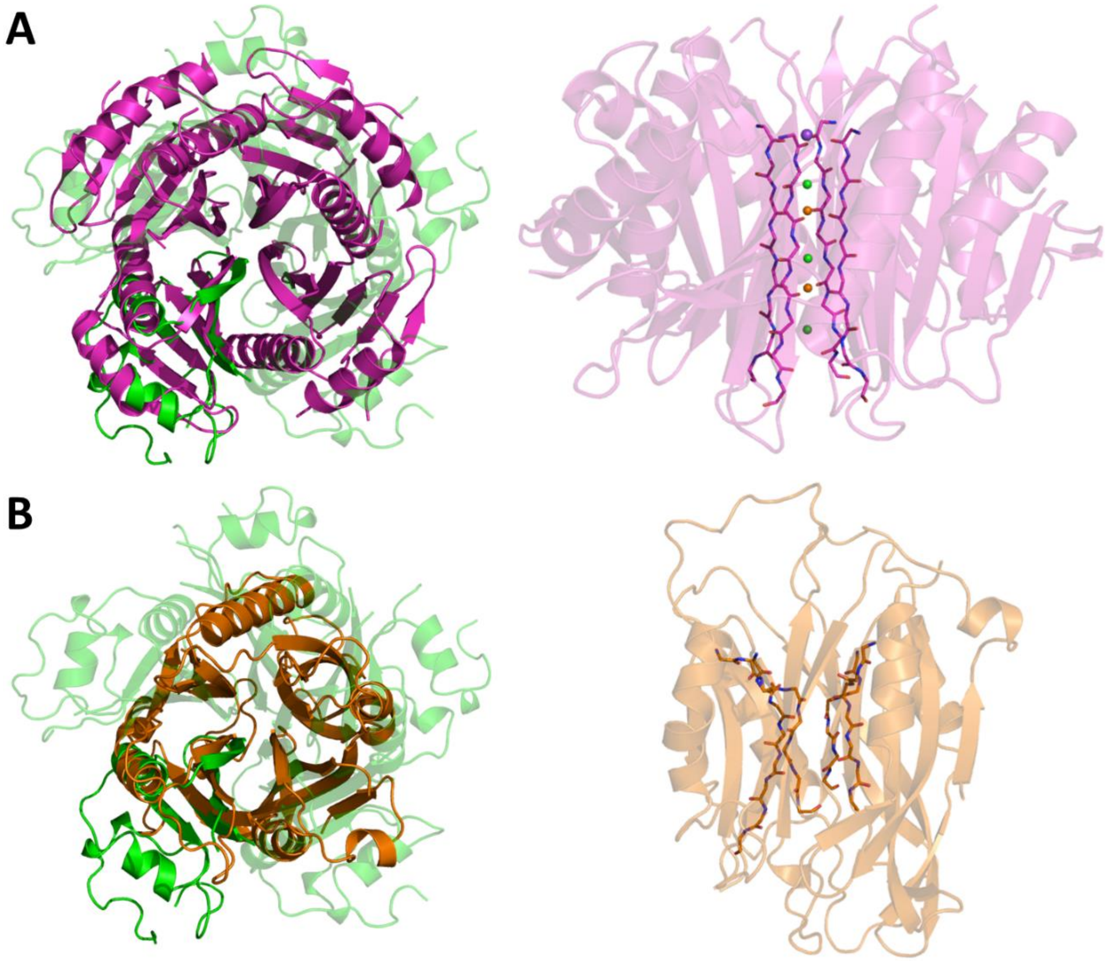


**Fig. S10.** Structural alignment of the CTD domain of KCTD1 (green) with the paraflagellar rod component (magenta, UniProtKB Q4D6Q6, PDB ID 6XYD) (A), and Raftlin-2 (orange, UniProtKB Q52LD8, residues 16-165 and 242-402 of the AF model) (B). Different views of these proteins with residues forming the central channel shown as sticks (only main chain atoms) are reported on the right. Magnesium (orange), sodium (purple), and chloride (green) ions inside the channel of the paraflagellar rod component are shown as spheres.

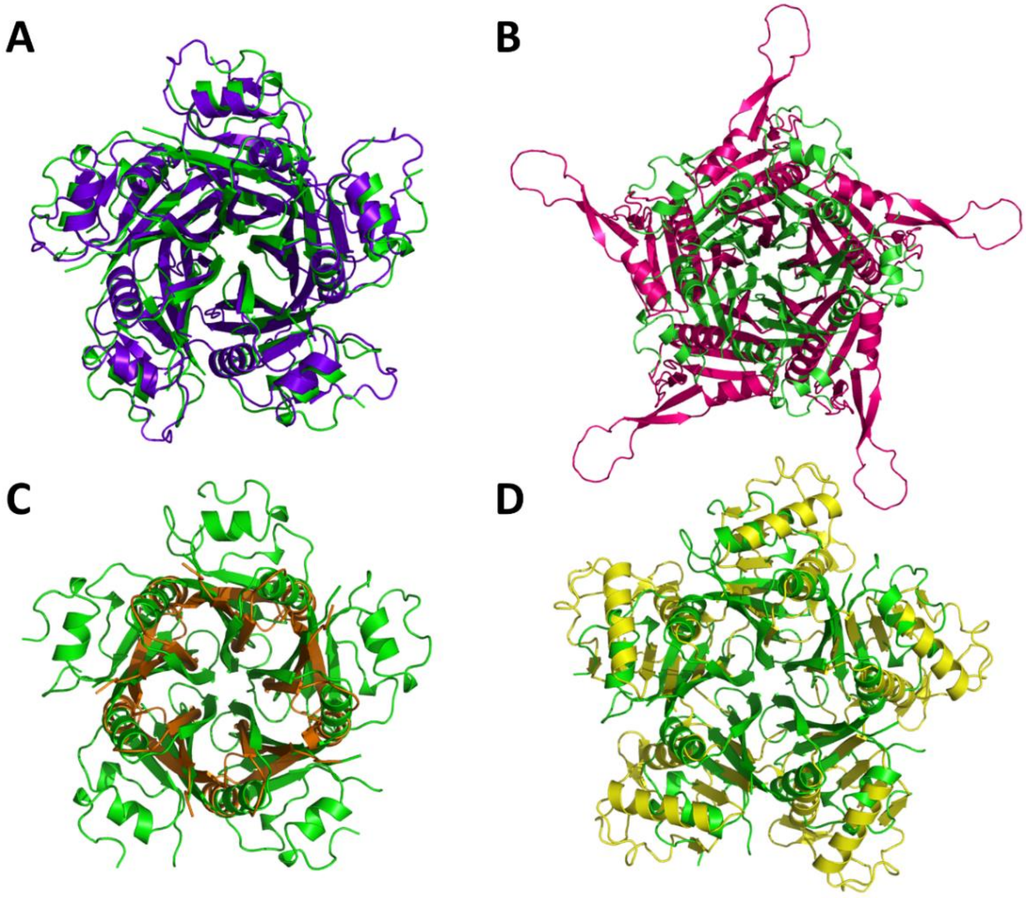


**Fig. S11.** Structural alignment of the CTD domains of KCTD1 (green) and (A) KCTD12 (blue), (B) KCNRG (magenta), (C) KCTD5 (orange), and (D) KCTD4 (yellow). The structural superimpositions were obtained by combining US-align and PyMOL.

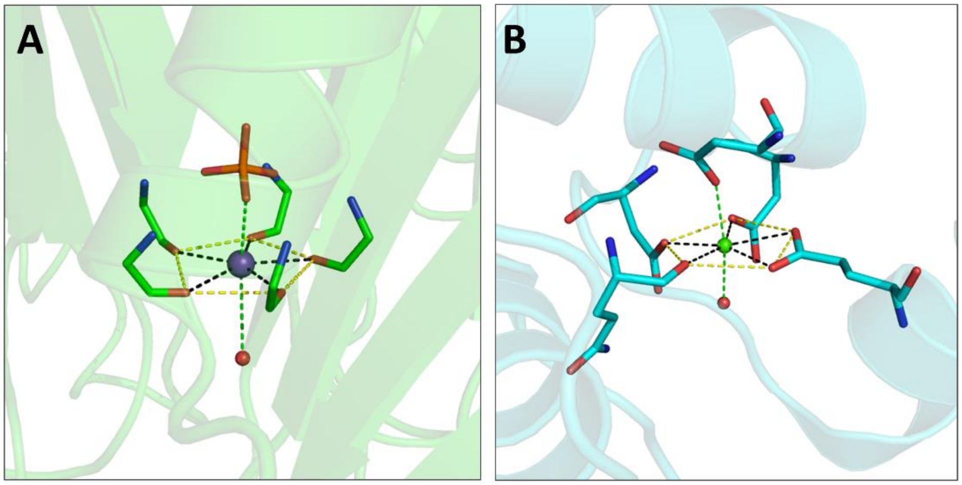


**Fig. S12.** Analogy of the metal coordination between KCTD1$^{P20S}$ and calmodulin (PDB ID 5A2H). The potassium ion (purple) in KCTD1, the calcium ion (green) in calmodulin and water molecules (red) are shown as spheres.

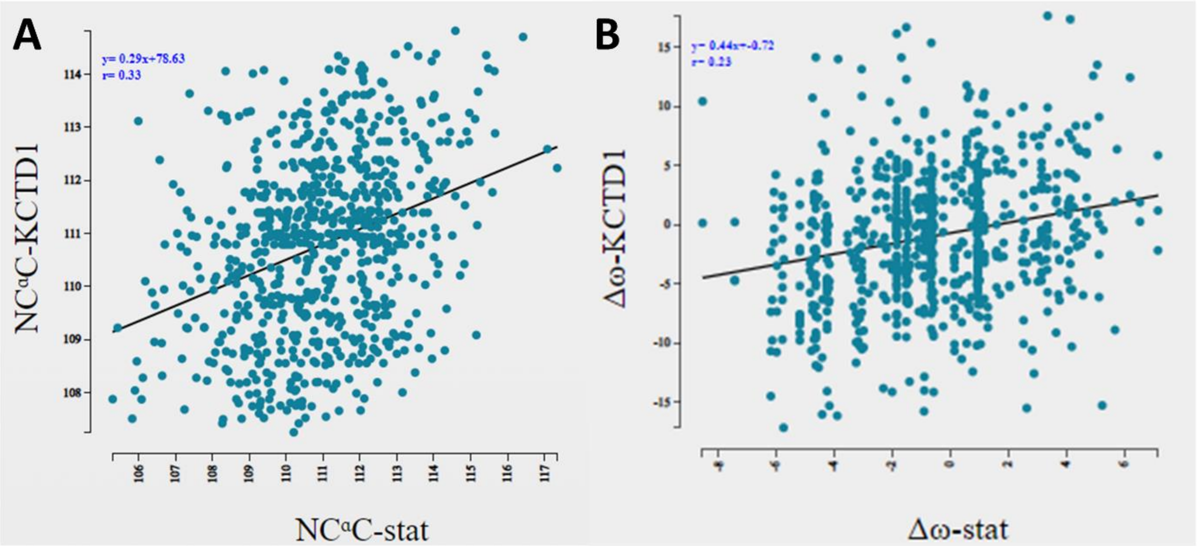


**Fig. S13.** Validation of the variability of some geometrical parameters of the protein backbone: the NC$^{\alpha}$C bond angle (A) and the deviation from the peptide bond planarity Δω defined as (ω-180°)mod360° (B) of KCTD1$^{P20S}$ crystal structure. The regression analysis has been performed by plotting the NC$^{\alpha}$C or Δω values of KCTD1$^{P20S}$ residues *versus* the average NC$^{\alpha}$C or Δω values of residues adopting the same (φ, ψ) conformation obtained from databases of X-ray protein structures solved at high resolution (resolution <1.6 Å for NC$^{\alpha}$C and resolution < 1.2 Å for Δω). The regression lines are shown.

**Table S1.** Percentages of sequence identity, computed on the different domains of KCTD1 (UniProKB Q719H9) and its long isoform (UniProKB A0A2U3U043), between the human protein and its orthologs isolated from five species: *Mus Musculus, Balaenoptera acutorostrata, Caretta caretta, Gallus gallus, and Danio rerio.*

| **KCTD1** | ***Mus Musculus*** | ***Balaenoptera acutorostrata*** | ***Caretta caretta*** | ***Gallus gallus*** | ***Danio rerio*** |
|---|---|---|---|---|---|
| | ***Percentage of sequence identity (%)*** | | | | |
| **PreBTB** (residues 1-29) | 100 | 100 | 100 | 100 | 62.1 |
| **BTB** (residues 30-133) | 100 | 100 | 97.1 | 96.2 | 87.5 |
| **CTD** (residues 139-239) | 100 | 100 | 99.0 | 99.0 | 90.1 |
| **PostCTD** (residues 240-257) | 100 | 94.4 | 88.9 | 44.4 | 65.0 |
| **Long isoform** (UniProKB A0A2U3U043) | 97.2 | 98.5 | 79.4 | 84.8 | 70.3 |

**Table S2.** Sizes of the funnel of KCTD proteins whose CTD is endowed with a propeller-like fold. For each plane in the funnel the residues and the distances between the carbonyl oxygen atoms (O-O distance) and between the oxygen atoms and the center (M) of the circumcircle of the regular pentagon (formed by the O-O distances) are reported.

| Cluster | Protein | Planes | O-O distance (Å) | M-O distance (Å) |
|---|---|---|---|---|
| 1A | KCTD8 | V291 | 5.5 | 4.7 |
| | | C293 | 4.3 | 3.7 |
| | | S295 | 3.6 | 3.1 |
| | | G297 | 3.8 | 3.2 |
| | KCTD12 | V292 | 5.3 | 4.5 |
| | | C294 | 4.2 | 3.6 |
| | | S296 | 4.1 | 3.5 |
| | | G298 | 4.4 | 3.7 |
| | KCTD16 | V248 | 5.6 | 4.8 |
| | | C250 | 4.4 | 3.7 |
| | | S252 | 4.8 | 4.1 |
| | | V254 | 6.1 | 5.2 |
| 1B | KCTD1 | S220 | 3.9 | 3.3 |
| | | G222 | 3.1 | 2.6 |
| | | G224 | 3.8 | 3.2 |
| | KCTD15 | S246 | 4.1 | 3.5 |
| | | G248 | 3.3 | 2.8 |
| | | G250 | 3.9 | 3.3 |
| 2A | KCNRG | V226 | 4.4 | 3.7 |
| | | T228 | 5.3 | 4.5 |
| | | T230 | 6.3 | 5.4 |
| | KCTD6 | R206 | 5.1 | 4.3 |
| | | T208 | 5.1 | 4.3 |
| | | V210 | 5.6 | 4.8 |
| 2B | KCTD11 | D251 | 8.1 | 6.9 |
| | KCTD21 | S240OG | 3.6 | 3.1 |
| 3 | KCTD2 | E211 | 5.9 | 5.0 |
| | | L213 | 6.3 | 5.4 |
| | | S215 | 7.1 | 6.0 |
| | KCTD5 | E182 | 6.0 | 5.1 |
| | | L184 | 6.3 | 5.4 |
| | | S186 | 6.9 | 5.9 |
| | KCTD17 | E168 | 6.0 | 5.1 |
| | | L170 | 6.3 | 5.4 |

| | | | | |
|---|---|---|---|---|
| | | N172 | 6.5 | 5.5 |
| 4 | KCTD4 | L239 | 7.6 | 6.5 |
| | | S241 | 7.9 | 6.7 |

**Table S3.** Results of the regression analysis of the geometrical parameters of KCTD1$^{P20S}$ compared to those found in high-resolution crystal structures (resolution <1.6 Å for bond angles and < 1.2 Å for dihedrals). The peptide bond deviations from planarity $\Delta\omega$ is defined as $(\omega - 180°) \mathrm{mod}\, 360°$ whereas the carbonyl carbon pyramidalization $\theta_C$ is defined as $(\omega - \omega 3 + 180°) \mathrm{mod}\, 360°$ ($\omega 3$ is the dihedral angle defined by the atoms $OCN_{+1}C^{\alpha}_{+1}$).

| **Angles (°)** | **Correlation coefficient R** | **p-value** |
|---|---|---|
| $\Delta\omega$ | 0.23 | $<10^{-6}$ |
| $\theta_C$ | -0.09 | 0.018 |
| $NC^{\alpha}C$ | 0.33 | $<10^{-6}$ |
| $NC^{\alpha}C^{\beta}$ | 0.11 | $2.9*10^{-3}$ |
| $C^{\beta}C^{\alpha}C$ | 0.14 | $10^{-4}$ |
| $C^{\alpha}CO$ | 0.18 | $2*10^{-6}$ |
| $C^{\alpha}CN^{+1}$ | 0.20 | $<10^{-6}$ |
| $OCN^{+1}$ | 0.06 | 0.093 |
| $C^{-1}NC^{\alpha}$ | 0.21 | $<10^{-6}$ |
